# When the psychedelic state’s over: limited evidence for persistent neurophysiological changes in naturalistic psychedelic users

**DOI:** 10.64898/2026.03.30.711922

**Authors:** Maja Wójcik, Paweł Orłowski, Stanisław Adamczyk, Paweł Lenartowicz, Justyna Hobot, Michał Wierzchoń, Michał Bola

## Abstract

**Background:** Contemporary research indicates that psychedelics induce notable neurophysiological changes, some lasting weeks to months after a single dose. However, most evidence derives from acute administration studies and limited post-acute follow-ups. Long-term naturalistic psychedelic users remain critically underexamined, yet may exhibit distinct neurobiological profiles informing our understanding of persistent alterations following repeated exposure.

**Methods:** We recorded resting-state EEG in 57 long-term psychedelic users (abstinent ≥30 days) and 49 matched non-users across two independent sites under eyes-open and eyes-closed conditions. We analyzed oscillatory power, signal complexity, and source-localized effective connectivity, focusing on five canonical frequency bands and regions of the Default Mode, Salience, and Central Executive Networks. Analyses included linear mixed-effects modeling for power spectra and complexity results and a rank-based approach combining ordinary least squares regression with randomization inference for effective connectivity.

**Results:** We observed predominantly null findings. No significant between-group differences emerged for oscillatory power. Complexity comparison yielded results contrary to our hypothesis: psychedelic users exhibited lower complexity values in the eyes-open condition. Effective connectivity revealed no within- or between-network differences that would survive statistical corrections. Additionally, we report a few small-magnitude effects uncovered by exploratory analyses.

Conclusions

Long-term naturalistic psychedelic users showed largely non-significant differences in oscillatory power, complexity, and network connectivity compared to non-users — across several measures commonly reported as altered in acute administration studies. These findings raise the question of whether psychedelics’ neurophysiological signatures persist during abstinence despite repeated prior use, or whether they reflect homeostatic receptor adaptation, individual variability, or contextual factors. Null, incongruous, or subtle effects contribute to the existing evidence base, yet underscore the need for replication in larger, more ecologically valid populations to advance the emerging field of psychedelic neuroscience.

## Introduction

In recent decades, psychedelic research has expanded rapidly, reframing substances like LSD (lysergic acid diethylamide) or psilocybin from criminalized drugs into investigational therapeutics (Aday et al., 2019). This “psychedelic renaissance” has been marked by three key developments: the Food and Drug Administration’s designation of certain psychedelic-assisted therapies as “breakthrough treatments” (Behera et al., 2024; Meling et al., 2024), the reshaping of drug policy across several countries, and the rapid expansion of scientific research on psychedelics (Andrews et al., 2025; Carhart-Harris et al., 2022; Johansen & Krebs, 2015).

### Acute effects and mechanistic models

Clinical neuroimaging studies using functional magnetic resonance imaging (fMRI), magneto-/electroencephalography (MEG/EEG), and a range of complementary measures such as magnetic resonance spectroscopy or positron emission tomography consistently report robust neurophysiological alterations during acute psychedelic administration (Aday et al., 2019). Many of these findings form the empirical foundation for prominent mechanistic models of psychedelic action involving therapeutic or psychological translations, such as the: Relaxed Beliefs Under Psychedelics Model (REBUS; Carhart-Harris & Friston, 2019), Entropic Brain Hypothesis (EBH; Carhart-Harris et al., 2014), Claustro-Cortical Circuit and Cortico-Striato-Thalamo-Cortical models (Barrett, Krimmel, et al., 2020; Gattuso et al., 2023; Preller et al., 2018; Tagliazucchi et al., 2014, 2016), neuroplasticity frameworks (Calder & Hasler, 2023; Grieco et al., 2022; Ly et al., 2018; Vollenweider & Kometer, 2010) or even the broader: Dynamical Systems/Repertoires (Atasoy et al., 2017; Lord et al., 2019; Roseman et al., 2014a; Tagliazucchi et al., 2014), Integrated-Information (Tononi, 2004) and Triple Network Theory (TNT; B. Menon, 2019; V. Menon, 2011; V. Menon & Uddin, 2010).

Investigations of acute psilocybin effects in healthy volunteers under resting-state conditions demonstrated widespread cortical desynchronization of neurons using fMRI (Siegel et al., 2024), also reflected as significant broadband reductions of oscillatory power when employing MEG/EEG (Kometer et al., 2015; Muthukumaraswamy et al., 2013). Besides spectra, the temporal structure of brain activity is frequently characterized and examined by quantifying complexity of EEG signals. In the context of neuroscience, employing complexity measures has the aim of calculating the variability inherent in neural signalling — the amount of incompressible information or the unpredictability of subsequent neural states. Such variability emerges from dynamic interactions between neurons, local circuits, and large-scale networks, extending across wide spatiotemporal scales in the brain (Keshmiri, 2020). Acute psychedelic states have been characterized by widespread increases in neural signal complexity, which is hypothesized to reflect an enhanced information-processing capacity and greater repertoires of accessible brain states (Atasoy et al., 2017; Carhart-Harris, 2018; Carhart-Harris et al., 2014; Lord et al., 2019; Roseman et al., 2014a; Schartner et al., 2017; Tagliazucchi et al., 2014). Yet, recent research points out important nuances. For example, McCulloch et al. (2023; preprint) evaluated 13 entropy measures in 28 healthy volunteers scanned during acute psilocybin influence, comparing their findings to prior studies. Only 5 of 13 metrics showed significant associations with plasma psilocybin levels, with just 2 replicating previous findings. Limited inter-measure correlations were described as suggesting these complexity metrics might index partially distinct aspects of neural dynamics rather than reflecting a single, unified “brain entropy” construct.

Neuroimaging studies with psychedelic administration also yield notable findings on distinctive connectivity patterns during acute states: reduced within-connectivity and integration of resting-state networks (RSN), especially substantial reductions of DMN connectivity, frequently linked with enhanced whole-brain coupling between diverse regions (Gattuso et al., 2023; Knudsen, 2023; McCulloch, Knudsen, et al., 2022; Roseman et al., 2014b; Stoliker et al., 2023). Such patterns reflect two key phenomena: *desynchronization* (less coordinated activity among neural populations), and *desegregation* (reduced within-network integration with consequent loss of modular organization across canonical RSNs), while between-network and whole-brain connectivity of distant regions increase (Gattuso et al., 2023; Madsen et al., 2021; Siegel et al., 2024; Tagliazucchi et al., 2014). Given their fundamental relationship to connectivity dynamics, desynchronization and desegregation should manifest detectably in oscillatory power and complexity measures, respectively.

### Lasting changes

Beyond acute effects, some evidence suggests psychedelics might also produce long-term neurobiological changes (Aday et al., 2020). Two studies using fMRI have documented lasting neurophysiological changes, which included: reduced DMN connectivity/integrity with enhanced “global network flexibility” or reduced segregation (i.e. increased functional connectivity between networks), lasting from one week to three months following single psilocybin administration (both n=10 healthy volunteers; Barrett et al., 2020; Madsen et al., 2020). Using a similar design, one trial did not find evidence of persisting changes (McCulloch, Madsen, et al., 2022). More recently, Siegel et al. (2024) reported a wide range of persistent alterations, including hippocampus-DMN functional connectivity reduction, decreased network segregation/desynchronization and other changes, persisting up to 3 weeks after a high dose of psilocybin in fMRI multi-repeated scanning. Jointly with Subramanian et al. (2025), a paper describing the above dataset, authors strongly suggest these results represent the new neuroanatomical and mechanistic correlates of psychedelics’ therapeutic and long-term effects. However, they also reported that: (1) whole-brain connectivity level did not change following psilocybin (analysis of: 1–21 days post-psilocybin to predrug baseline); (2) hippocampus-DMN connectivity returned to pre-psilocybin baseline by the replication visit 6–12 months later; (3) that psilocybin acutely produced context-dependent desynchronization of brain activity and performing perceptual tasks reduced psilocybin-driven functional connectivity changes.

Cross-sectional studies on chronic ayahuasca users further reported cortical thickness differences (posterior cingulate cortex thickening) alongside personality changes correlating with onset and lifetime ayahuasca use (Bouso et al., 2015; n=22 per group; active church members vs. matched controls). More recently, Ramos et al. (2025) demonstrated that machine-learning algorithms correctly classified long-term ceremonial ayahuasca users (n=18 Brazilian males) based on fMRI patterns and predicted individual psychological resilience scores, although conventional univariate statistical comparisons revealed no between-group differences.

Both the acute and post-acute neurobiological changes of brain dynamics are frequently proposed as mechanistic explanations underlying treatment efficacy in depression (Daws et al., 2022; Thomas et al., 2017; Vollenweider & Kometer, 2010), anxiety (Bouso et al., 2021), PTSD, addiction and various other psychiatric disorders (Gattuso et al., 2023; Grieco et al., 2022; Schindler et al., 2018); in addition to mood elevation or psychological and personality traits reshaping (Aday et al., 2020; Barrett, Doss, et al., 2020; Bouso et al., 2021; Dos Santos et al., 2016; Knudsen, 2023; Lebedev et al., 2016). However, it still remains an open question whether those lasting beneficial effects derive solely from the acute psychedelic action or from any enduring alterations of neural circuit dynamics. It has been extensively demonstrated that psychological effects persist after subjective effects resolve — but whether neurobiological signatures last still needs to be established (Aday et al., 2020; Carhart-Harris, 2018; Gattuso et al., 2023; Grieco et al., 2022; Knudsen, 2023). Current evidence regarding long-term neurobiological changes remains limited by lack of longitudinal research and small, non-generalizable samples (Aday et al., 2020; Gattuso et al., 2023).

### Naturalistic users

While chronic ayahuasca users have received some research attention, those people often remain culturally/contextually distinct from other groups of naturalistic psychedelic users. Drawing predominantly on male participants rooted in religious or indigenous traditions and ceremonial use (Dos Santos et al., 2016), studies of ayahuasca users contrast sharply with, for example, more WEIRD (Western, Educated, Industrialized, Rich, Democratic) populations that make up the majority of psilocybin/LSD studies (Aday et al., 2020; George et al., 2022; Michaels et al., 2018). Controlled clinical trials have rigorously documented acute administration effects, but our knowledge regarding neurophysiological characteristics of people using psychedelics on their own, naturalistically — outside of laboratory or clinical context — remains critically incomplete (Aday et al., 2019; Schenberg, 2024). Although essential, clinical trials do not fully capture the complexity of factors largely influencing long-term outcomes profiles. Setting, intention, integration practices, baseline state or inter-individual variability are largely expected to impact the outcomes but remain poorly examined (Adamczyk et al., 2025; Aday et al., 2020; Carhart-Harris, 2018; Hartogsohn, 2016, 2016; Michaels et al., 2018; Nutt et al., 2020; Strassman, 1991). Examining ecologically valid populations — with greater diversity in individual characteristics, dosage patterns, environmental and cultural contexts — seems therefore essential to fill this remaining research gap. Studies on cumulative and prolonged use, boundary conditions could advance the existing mechanistic theories (Aday et al., 2020; Carhart-Harris et al., 2022; Michaels et al., 2018; Schenberg, 2024) .

There are emerging studies that have examined psychedelic-induced acute neural effects - occurring in more naturalistic conditions incorporating in laboratory-based settings that were individually preferred by participants (i.e. music). Employing this design, Pallavicini et al. (2021) identified alterations of spectral power — decreased alpha and increased gamma and delta — and marked as potential neural correlates of mystical experiences in healthy, experienced participants, following DMT inhalation. Similarly, Tagliazucchi et al. (2021) found that DMT acutely altered spectral dynamics in psychedelic-naive participants, which correlated with aspects of conscious experience. Beyond these acute investigations, our prior analysis revealed altered emotional processing, including reduced early-stage reactivity to negatively-valenced emotional stimuli in long-term naturalistic users, abstinent during study (Orłowski et al., 2024b).

### Current study

Building on this evidence, our aim was to investigate whether long-term naturalistic psychedelic users would exhibit neural signatures similar to those observed in acute and post-acute states - in comparison to matched non-users. Given limited research on repeated or naturalistic psychedelic use, we based our hypotheses on reports from several significant clinical studies and assumptions of selected mechanistic models. We employed three complementary resting-state EEG signal analyses: oscillatory power via power spectrum density (PSD), signal complexity via the Lempel-Ziv algorithm (LZ), and effective connectivity between regions of interest (ROIs) of DMN, Salience Network (SN), and Central Executive Network (CEN) via source-localized non-normalized Directed Transfer Function (nDTF; Kaminski & Blinowska, 1991). PSD and effective connectivity were analysed across five frequency bands: delta (1–3 Hz), theta (4–8 Hz), alpha (9–13 Hz), beta (14–30 Hz), and gamma (31–45 Hz).

We hypothesized that long-term naturalistic psychedelic users, compared to non-users, would demonstrate: (1) lower global oscillatory power, particularly in alpha and beta frequency bands (Kometer et al., 2015; Muthukumaraswamy et al., 2013; Ort et al., 2023; Schartner et al., 2017; Vejmola et al., 2021) (2) higher neural signal complexity (Carhart-Harris, 2018; Carhart-Harris et al., 2014; Murray et al., 2024; Schartner et al., 2017; Timmermann et al., 2019, 2019); (3) lower effective connectivity within DMN-DMN and SN-SN regions; alongside (4) higher between-network effective connectivity, particularly within ROIs comprising DMN-to-CEN and SN-to-DMN connections (Bouso et al., 2015; Castelhano et al., 2021; Gattuso et al., 2023; Madsen et al., 2021; Roseman et al., 2014a; Stoliker et al., 2021, 2023; Tagliazucchi et al., 2016); and (5) condition-specific differences between eyes-open and eyes-closed states (Agcaoglu et al., 2019; McCulloch, Knudsen, et al., 2022). Hypotheses 1-3 were preregistered (see: Hobot et al., 2021; https://osf.io/z6tky). Hypotheses 4-5 were added based on additional literature review (after the start of data collection) and should be considered exploratory.

## Methods

This research was conducted in accordance with the Declaration of Helsinki and approved by the Human Ethics Committee of SWPS University of Social Sciences and Humanities (Warszawa, Poland; approval 13/2020) and the Jagiellonian University Bioethics Committee (Kraków, Poland; approval 1072.6120.259.2021). Data reported here are a part of a larger project, with some results already published: Adamczyk et al., 2025; Orłowski et al., 2022, 2024a, 2024b; and Ruban et al., 2025.

### Participants

We analyzed data from 57 long-term psychedelic users (M = 27.8 years, SD = 5.7; 21 males, 36 females) and 49 matched non-users (M = 28.6 years, SD = 5.8; 20 males, 29 females) collected across two independent sites: dataset I from Jagiellonian University (Kraków; n = 40; 21 psychedelic users) and dataset II from SWPS University (Warszawa; n = 66; 36 psychedelic users), following recommendations for independent replication in psychedelic research (McCulloch, Knudsen, et al., 2022).

We attempted both formal a priori sample size calculations during study design (Hobot et al., 2021) and a post-hoc power analysis; however, the scarcity of studies examining naturalistic users with comparable methodology rendered such estimates unreliable. We therefore recruited the largest practicable sample given available resources. For context, McCulloch et al. (2022) noted that studies evaluating psilocybin-induced changes in resting-state functional connectivity would require samples exceeding 60 participants to achieve adequate statistical power (i.e. 1−*β* > 0.8), given that most observed effect sizes for within- and between-network connectivity were small to medium (|Cohen’s *d*| < 0.5).

Participants were recruited through an online survey (LimeSurvey GmbH, v3.22.7) distributed primarily via Polish harm reduction organizations (politykanarkotykowa.pl; sin.org.pl; psychodeliki.org). The survey received approximately 5,000 responses (for details see Orłowski et al., 2022) and collected information on psychoactive substance history, sociodemographic characteristics, medication use, meditation practices, psychiatric/neurological history, and the abbreviated Alcohol Use Disorders Identification Test (AUDIT; items 1-4; Bush et al., 1998). An overview of recruitment, screening steps and included participants as a CONSORT-like flowchart is presented in Figure 1.

**Figure 1.**
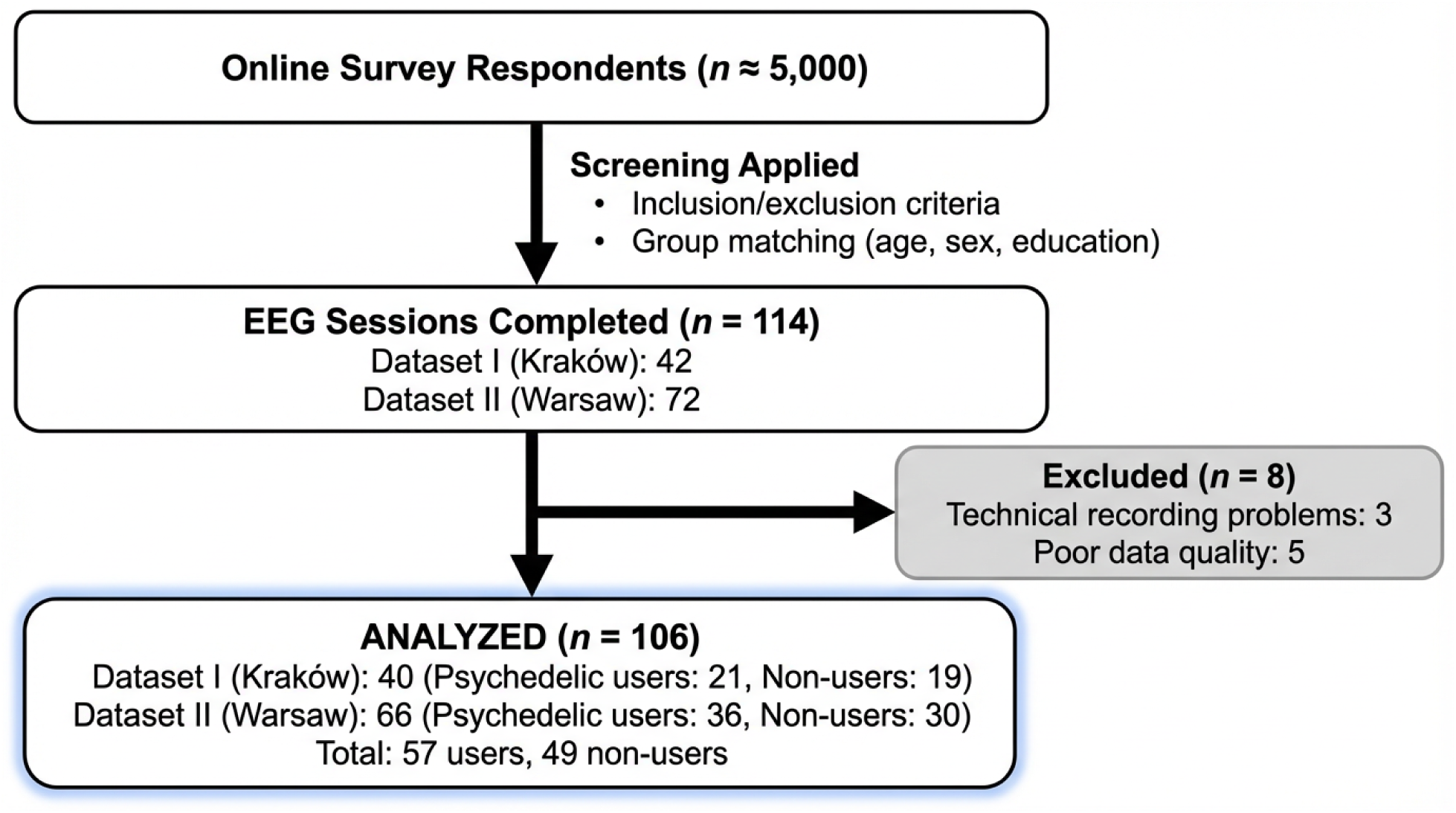
Participant Selection and Screening Overview in a CONSORT-like Flowchart.

Exclusion criteria were: frequent use of empathogens, stimulants or dissociatives (>15 lifetime or >5 past-year); any use of benzodiazepines, synthetic cannabinoids, or opioids (>3 lifetime or >1 past-year); AUDIT scores ≥4 on any item; or psychiatric/neurological diagnoses.

Psychedelic users group reported ≥15 lifetime experiences (M = 30.5, SD = 25.4) using psychedelics (LSD, psilocybin, mescaline, DMT/ayahuasca, and closely related synthetic derivatives such as 5-MeO-DMT and 4-AcO-DMT), explicitly excluding MDMA and ketamine. Participants were instructed to exclude microdosing and count only experiences involving significant perceptual alterations at standard doses. The non-users group had never tried psychedelics but expressed willingness to do so in the future — a criterion enhancing group matching by controlling for curiosity/openness trait (increasingly recognized as one of the key features of long-term psychedelic effects; e.g. Aday et al., 2019; Knudsen, 2023).

The psychedelic users group were instructed to abstain from psychedelics for at least 30 days before the EEG session. Hair samples were collected from consenting participants on the study day; all participants were notified in advance that samples would undergo drug testing (described as a precaution to verify declared abstinence).

### Laboratory sessions

Core procedural steps were identical across all sessions. They started with the participant completing a set of three questionnaires: the Reflection-Rumination Questionnaire (RRQ; Takano & Tanno, 2009), State-Trait Anxiety Inventory (STAI; Spielberger, 2012), and Beck Depression Inventory-II (BDI-II; Beck et al., 2011). This was followed by a 10-minute resting-state EEG recording with two conditions (5 min eyes-closed; 5 min eyes-open). Directly after the resting-state recording, the participant filled the Amsterdam Resting-State Questionnaire (ARSQ; Diaz et al., 2013). Sessions concluded with completing three experimental tasks, not analyzed in the present study: emotional facial expression perception (Orłowski et al., 2024b), self-referential stimuli perception (Orłowski et al., 2024a), and rumination induction tasks (Ruban et al., 2025). More detailed description of the laboratory sessions is available in Supplementary Material 1.

#### EEG Recording

EEG signals were recorded from 64 scalp sites using Ag/AgCl active electrodes positioned according to the extended 10-20 system. Dataset I employed a BioSemi ActiveTwo amplifier system (BioSemi, Amsterdam, Netherlands) with BioSemi ActiView software at a 2048 Hz sampling rate. Dataset II utilized a Brain Products ActiCap system with Brain Vision Recorder software at a 1000 Hz sampling rate.

Detailed preprocessing pipelines are provided in Supplementary Material 2. All scripts, raw behavioral and EEG data are available in the study’s repository. EEG data underwent preprocessing via two parallel pipelines (maintained for maximal consistency): a standard pipeline for PSD and LZ analyses, and a more stringent, purpose-optimized pipeline for effective connectivity calculations to ensure robust statistical inference.

#### Oscillatory power

Oscillatory power analysis was conducted using a Python script incorporating MNE library functions for EEG processing (Gramfort et al., 2014). PSD was estimated using Welch’s method, which segments continuous signals into 4-second epochs. PSD was computed for each channel and epoch using MNE’s compute_psd() function, then averaged across all epochs and channels. The resulting power spectrum was partitioned into: delta, theta, alpha, beta, and gamma frequency bands. For each band, PSD values within the corresponding frequency range were averaged.PSD data were converted from V²/Hz to μV²/Hz, then log-transformed using the formula 10 × log₁₀(x), where x represents the input signal, yielding decibels microVolts²/Hertz (dB μV²/Hz). This transformation normalized the data distribution, following standard EEG preprocessing practices (Cohen, 2014; Delorme & Makeig, 2004).

#### Complexity

Signal complexity was assessed using the Lempel-Ziv (LZ) algorithm to evaluate signal diversity within individual EEG channels, following the algorithmic implementation of Schartner et al. (2017). For each channel, 4-second epochs of EEG data were processed independently. First, signals were demeaned and normalized by dividing by their standard deviation. Linear trends were subsequently removed, and signal envelopes were calculated using the Hilbert Transform. Signals were then binarized based on the mean value of their envelope, which served as the binarization threshold. The binary signal was analyzed using the Lempel-Ziv compression algorithm, which estimates signal diversity by counting unique patterns in the binary sequence. To ensure that the final values reflected intrinsic signal complexity rather than random fluctuations, raw LZ scores were normalized by dividing by LZ scores obtained from the same binary signal shuffled in time. The resulting normalized LZ values ranged from 0 (minimal diversity) to 1 (maximal diversity). This normalization controls for differences in sequence length and ensures comparability across channels and participants.

#### Effective Connectivity

Effective connectivity was estimated across delta, theta, alpha, beta, and gamma frequency bands for three resting-state networks: DMN, CEN, and SN, comprising a total of 22 ROIs. ROI selection was guided by the Triple Network Theory (TNT; V. Menon, 2011) and partly the Raichle’s RSNs parcellation (Raichle, 2011; Raichle et al., 2001), frameworks consistently implicated in both psychedelic research (e.g. Barrett, Krimmel, et al., 2020; Gattuso et al., 2023; McCulloch, Madsen, et al., 2022; Roseman et al., 2014) and significant cognitive and clinical neuroscience studies (e.g. Andrews-Hanna et al., 2007; Putcha et al., 2016; Sherman et al., 2014a; Smallwood et al., 2021; Tops et al., 2014).

ROI placement was determined through convergent evidence from neuroanatomical atlases (Harvard-Oxford Cortical Structure Atlas) and literature (Agcaoglu et al., 2019; Pietrzykowski et al., 2022; Seeley et al., 2007; Shao et al., 2018; Sherman et al., 2014b; Smallwood et al., 2021; Tops et al., 2014; Wang et al., 2018). To ensure robust source reconstruction given the EEG methods inverse problem (Grech et al., 2008; Pascual-Marqui, 2007), ROIs were required to: (1) maintain minimum 20mm inter-ROI distance to minimize spatial leakage, (2) encompass ≥70% cortical tissue, which was verified using FSLeyes (Jenkinson et al., 2012), and (3) overlap ≥40% with anatomical labels in the Harvard-Oxford atlas on the MNI152_T1_1mm brain template. These constraints balanced anatomical fidelity with computational tractability, ensuring the 22-ROI model remained appropriate for inverse solution algorithms while preserving functionally meaningful network architecture. 22 ROIs of our selection are visualised in Figure 2 (BrainNet Viewer, Xia et al., 2013). Exact MNI coordinates of the ROIs are provided in Supplementary Material 3, Table S2.

**Figure 2.**
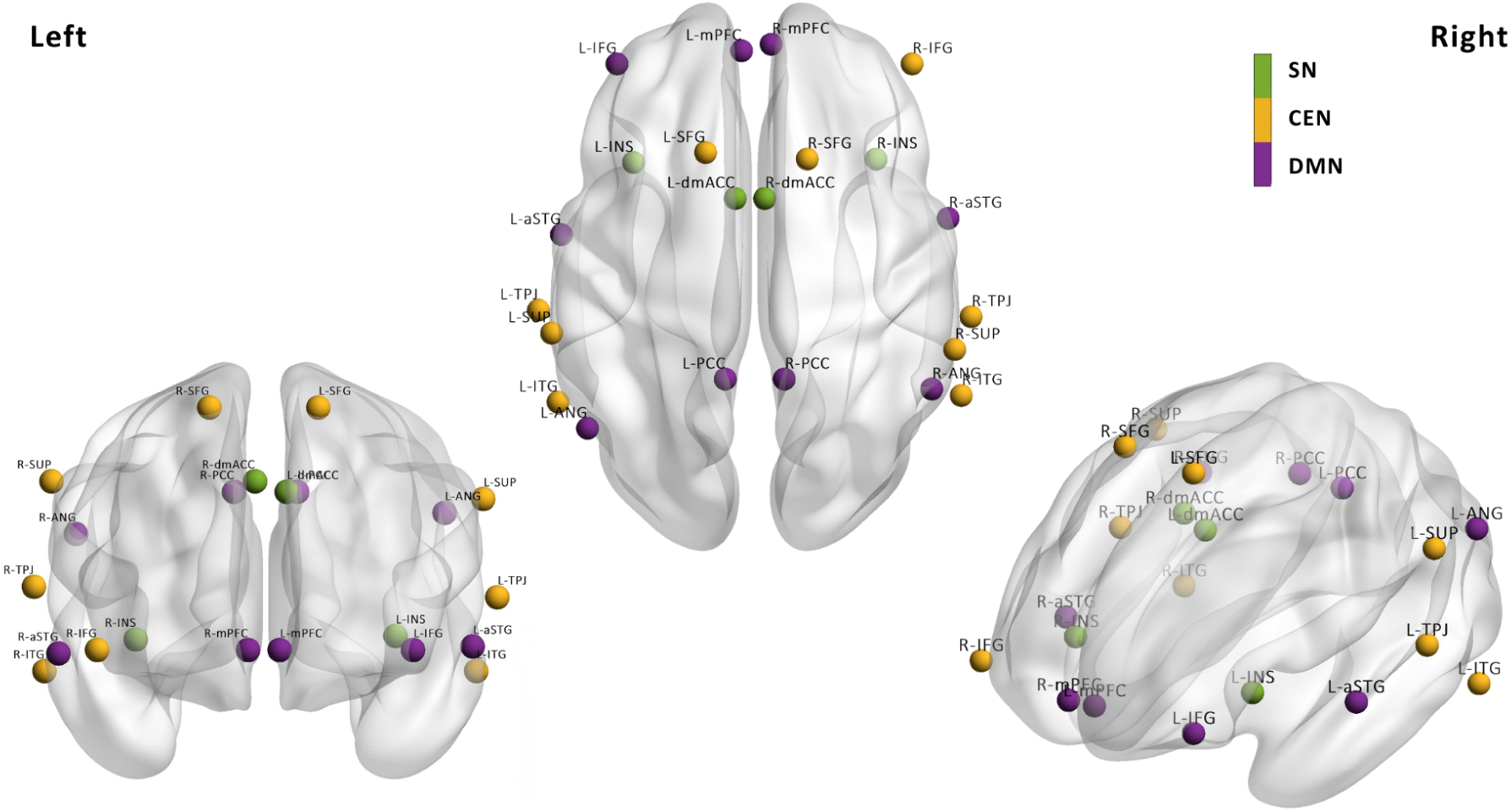
Spatial Visualisation of 22 Selected Regions of Interest By Three Networks. This visualisation was created using BrainNet Viewer (Xia et al., 2013). SN = Salience Network; DMN = Default Mode Network; CEN = Central Executive Network; dmACC = dorso-medial anterior cingulate cortex; INS = insular cortex; IFG = inferior frontal gyrus; mPFC = medial prefrontal cortex; aSTG = anterior superior temporal gyrus; TPJ = temporal parietal junction; PREC – precuneus; PCC = posterior cingulate cortex; ANG = angular gyrus; SFG = superior frontal gyrus; pSTG = posterior superior temporal gyrus; ITG = inferior temporal gyrus; SUP = supramarginal gyrus.

Effective connectivity was estimated using non-normalised Directed Transfer Function (nDTF), a multivariate extension of Granger causality (Granger, 1969) that simultaneously quantifies directional information flow across all signals in the system (Blinowska, 2011; Kaminski & Blinowska, 1991). Unlike pairwise approaches, nDTF’s multivariate autoregressive modelling (MVAR) accounts for both direct and indirect influences among all specified ROIS, providing comprehensive characterization of directed functional interactions of regions. nDTF has been validated for EEG-based effective connectivity analyses and has been shown to be particularly suited for examining network-level dynamics in source space (for methodological reviews see: Bakhshayesh et al., 2019; Bastos & Schoffelen, 2016; Blinowska, 2011; Fakhar et al., 2025; Laasch et al., 2025; Winterhalder et al., 2005).

Computation of nDTF results was performed using the ASCT toolbox (https://asct.gitbook.io/asct-manual/; Wyczesany et al., 2025), with the analysis steps detailed in Supplementary Material 2. Complete MATlab pipelines employed for this study are available in the repository.

### Statistical analysis

#### Oscillatory power and Complexity

Linear mixed-effects models were used to assess the effects of group (psychedelic users vs. non-users), condition (eyes-open vs. eyes-closed), and their interaction on PSD and LZ. Data collection site was included as a fixed effect and participant as a random effect to account for between-site variability and within-participant dependencies. Post-hoc comparisons were conducted on estimated marginal means (Searle et al., 1980) using the emmeans R package (Lenth & Piaskowski, 2017), with p-values adjusted for multiple comparisons via the Holm-Bonferroni method. Additional analyses examining potential site effects are included in Supplementary Material 4, Table S3.

#### Effective connectivity

The nDTF dataset comprised measurements across groups (psychedelic users vs. non-users), conditions (eyes-open vs. eyes-closed), frequency bands (delta, theta, alpha, beta, gamma), and 462 directed ROI-to-ROI pairs (22 ROIs × 21 targets), yielding 4,620 unique data points (nDTF values) per participant (462 pairs × 5 bands × 2 conditions).

Given the inherently skewed distribution of nDTF data, we implemented a rank-based analytical approach to ensure robust statistical inference. Choosing a non-parametric strategy addressed the non-normality of nDTF values while preserving the relative ordering of connectivity strengths across participants.

For each of the 4,620 nDTF measurements, we calculated participant ranks within each condition after excluding missing values. Normalized Mann-Whitney U statistics were then computed for between-group comparisons. This approach provided a standardized effect size measure quantifying the magnitude of group differences while remaining robust to outliers and distributional assumptions.

ROI-to-ROI pairs were grouped according to their source and destination networks (see Supplementary Material 3), creating broader (network-to-network) categories (e.g., DMN-to-SN, CEN-to-CEN) for conducting network-based connectivity analyses.

We employed ordinary least squares (OLS) regression modeling with normalized Mann-Whitney U values (contrasts) as the dependent variable and three independent variables: condition, frequency band, and the network-based categories. Interaction terms between network categories and frequency bands were additionally included for capturing potential frequency-specific network effects.

Statistical significance was determined via randomization inference with 10,000 permutations of group assignments while maintaining all other data structure intact. Confidence intervals were derived from permutation-based standard errors.

All above analyses were implemented using custom Python code, included in the repository with comprehensive documentation, example datasets and the aim to ensure reproducibility and support further methodological extensions.

## Results

### Participant characteristics

Groups were well-matched on demographic characteristics across both datasets. No significant differences emerged in age, sex distribution, or education level (all *p*s > .05). All participants were native Polish speakers; most holding bachelor’s degrees or higher (dataset I: 76% users, 74% non-users; dataset II: 87% users, 85% non-users). For complete demographic data and statistical comparisons, see Table S1 in Supplementary Material 1.

Lifetime cannabis use did not differ significantly in dataset I (*p* = .544) but was higher among psychedelic users in dataset II (*p* = .004). Alcohol consumption (AUDIT scores) and lifetime meditation hours were comparable between groups in both datasets (all *p*s > .05). Psychological assessments including anxiety (STAI), depression (BDI-II), reflection-rumination (RRQ), and resting-state experience (ARSQ) measures revealed largely similar profiles between groups across both datasets. Although significant (small-magnitude) differences emerged for isolated subscales, their interpretation was deemed beyond this study’s scope (full results are provided in Supplementary Material 1, Figures S1–S8).

### Oscillatory power

No significant main effects of the group were observed in any frequency band, although statistically significant main effects of the condition were found across all frequency bands (Table 1). Specifically, delta, theta, alpha, and beta showed higher PSD values in the eyes-closed condition, while gamma showed higher PSD values in the eyes-open condition.

**Table 1.**
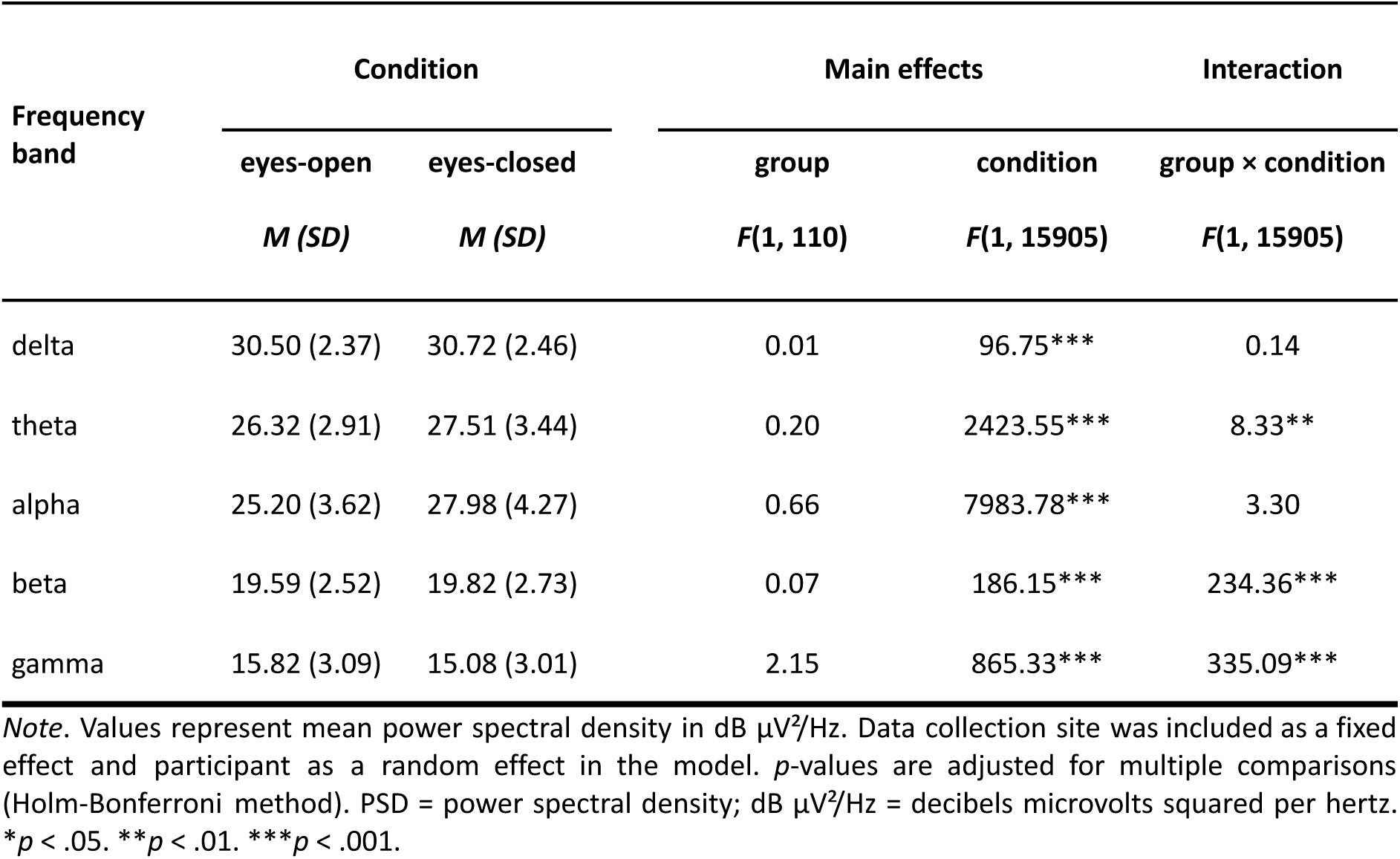
Results of Linear Mixed-Effects Model Analyses for Oscillatory Power Across All Frequency Bands.

| Frequency band | Condition |  | Main effects |  | Interaction |
| --- | --- | --- | --- | --- | --- |
|  | eyes-open | eyes-closed | group | condition | group × condition |
|  | *M (SD)* | *M (SD)* | *F*(1, 110) | *F*(1, 15905) | *F*(1, 15905) |
| delta | 30.50 (2.37) | 30.72 (2.46) | 0.01 | 96.75*** | 0.14 |
| theta | 26.32 (2.91) | 27.51 (3.44) | 0.20 | 2423.55*** | 8.33** |
| alpha | 25.20 (3.62) | 27.98 (4.27) | 0.66 | 7983.78*** | 3.30 |
| beta | 19.59 (2.52) | 19.82 (2.73) | 0.07 | 186.15*** | 234.36*** |
| gamma | 15.82 (3.09) | 15.08 (3.01) | 2.15 | 865.33*** | 335.09*** |

*Note.* Values represent mean power spectral density in dB $\mu\text{V}^2/\text{Hz}$ . Data collection site was included as a fixed effect and participant as a random effect in the model. *p*-values are adjusted for multiple comparisons (Holm-Bonferroni method). PSD = power spectral density; dB $\mu\text{V}^2/\text{Hz}$ = decibels microvolts squared per hertz. \**p* < .05. \*\**p* < .01. \*\*\**p* < .001.

Statistically significant group×condition interactions were observed for theta, beta, and gamma. Post-hoc analyses indicated that psychedelic users showed altered eyes-open vs. eyes-closed contrasts, compared to non-users, though the pattern of effects was inconsistent. In beta, psychedelic users demonstrated more pronounced between-condition contrast (of negative valence) compared to non-users. Conversely, in gamma psychedelic users exhibited smaller eyes-closed vs. eyes-open contrast compared to non-users (see Figure S9 in Supplementary Material 4 for full results). Lastly, a single significant between-group difference emerged in gamma, but only in the eyes-open condition, where non-users showed higher PSD values (Table 2).

**Table 2.**
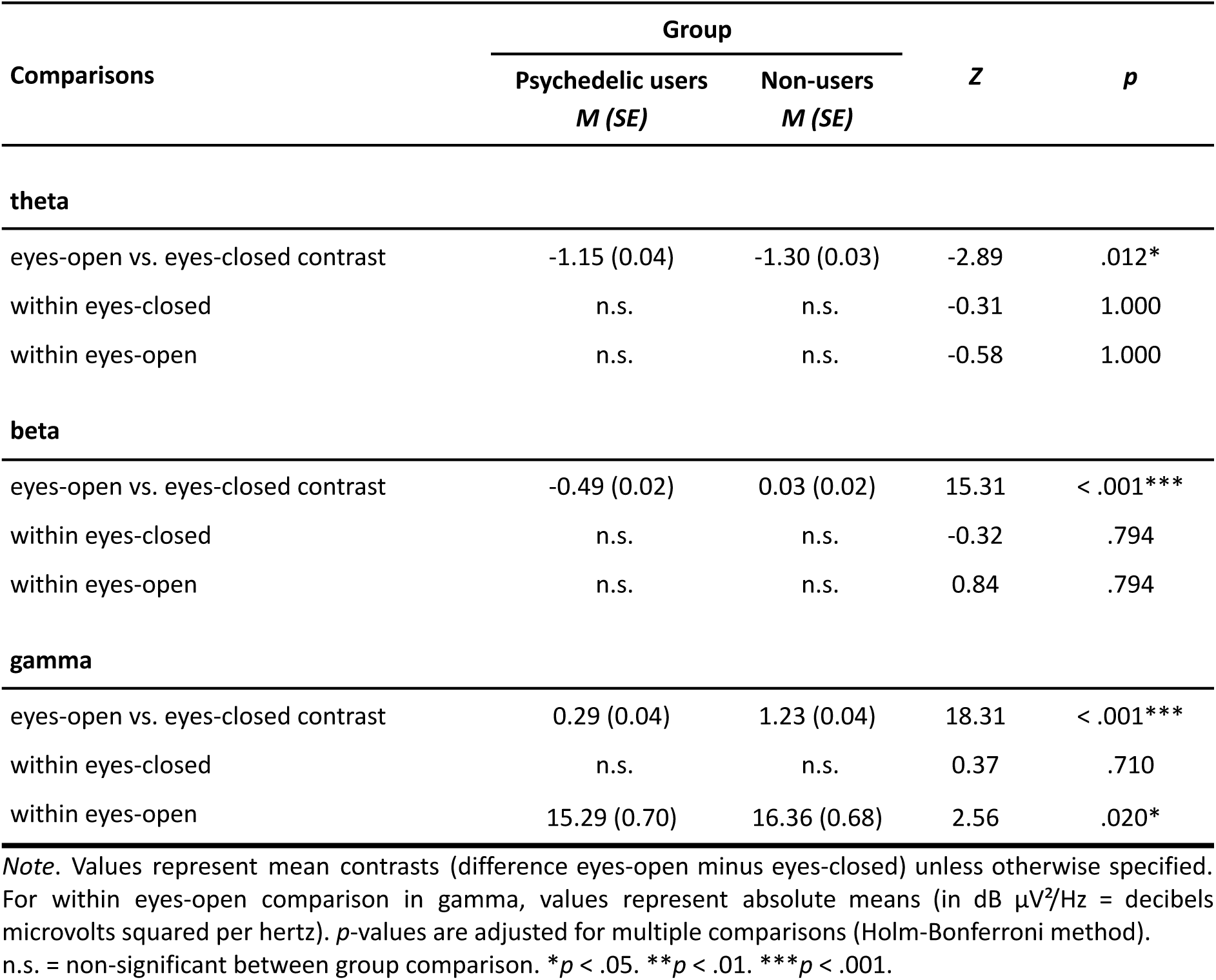
Post-Hoc Comparisons for Frequency Bands With Significant Group × Condition Interaction Effects.

### Complexity

No statistically significant main effect of the group was observed, *F*(1, 110) = 2.24, *p* = .137. However, a significant main effect of the condition emerged *F*(1, 15909) = 5685.42, *p* < .001, with higher LZ values in the eyes-open compared to eyes-closed condition. The group × condition interaction was also statistically significant *F*(1, 15909) = 141.24, *p* < .001. Post-hoc analyses revealed that psychedelic users group exhibited lower LZ values than the non-users in the eyes-open condition (*t* = 2.33, *p* = .022), contrary to our hypothesis. No significant between-group difference was detected in the eyes-closed condition (*t* = 0.66, *p* = .511; Figure 3).

**Figure 3.**
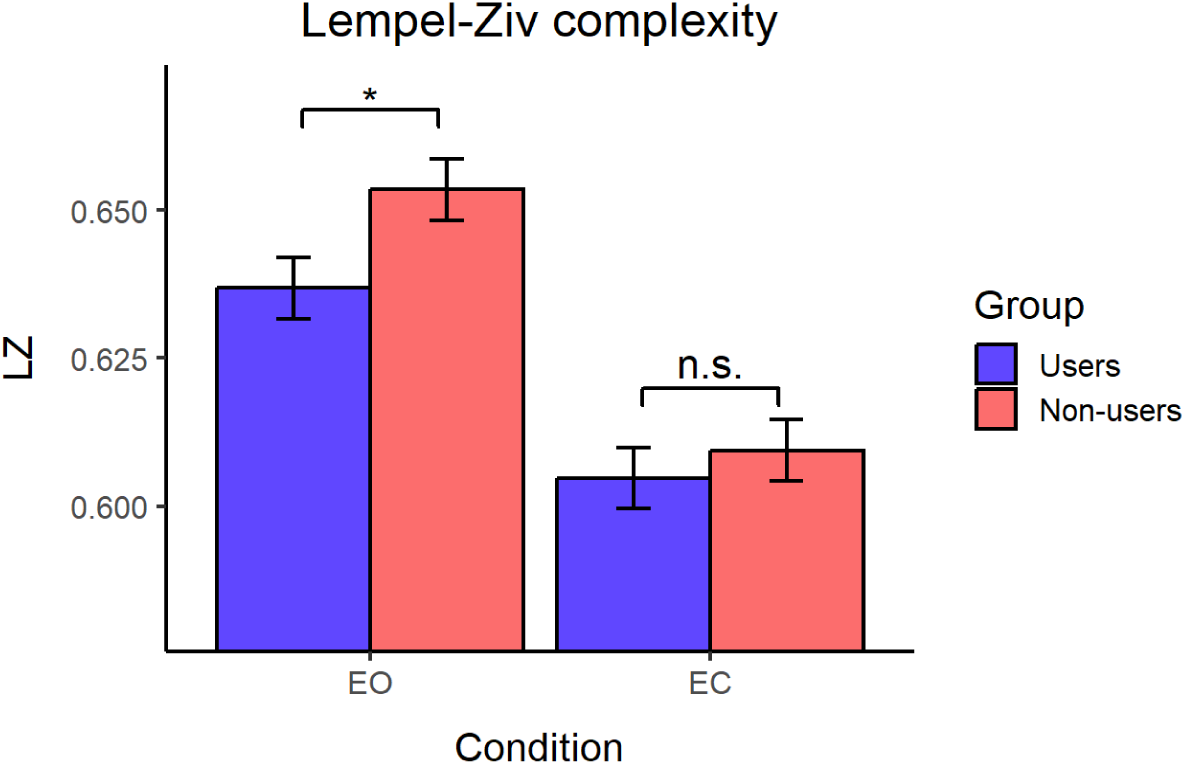
Lempel-Ziv Complexity Values by Condition and Group. Mean LZ complexity values for eyes-closed and eyes-open conditions, shown separately for psychedelic users and non-users. Error bars indicate standard error. LZ = Lempel-Ziv mean values; EO = eyes-open condition; EC = eyes-closed condition. *n.s.* = non-significant. *p* < .05*\**.

### Effective connectivity

No significant group effects were observed (Table 3). Contrary to hypothesis (1), psychedelic users did not exhibit altered patterns of effective connectivity within SN-SN or DMN-DMN regions. Contrary to hypothesis (2), DMN-to-CEN connection showed a small negative but non-significant effect, while SN-to-DMN exhibited minimal and also non-significant effect.

**Table 3.**
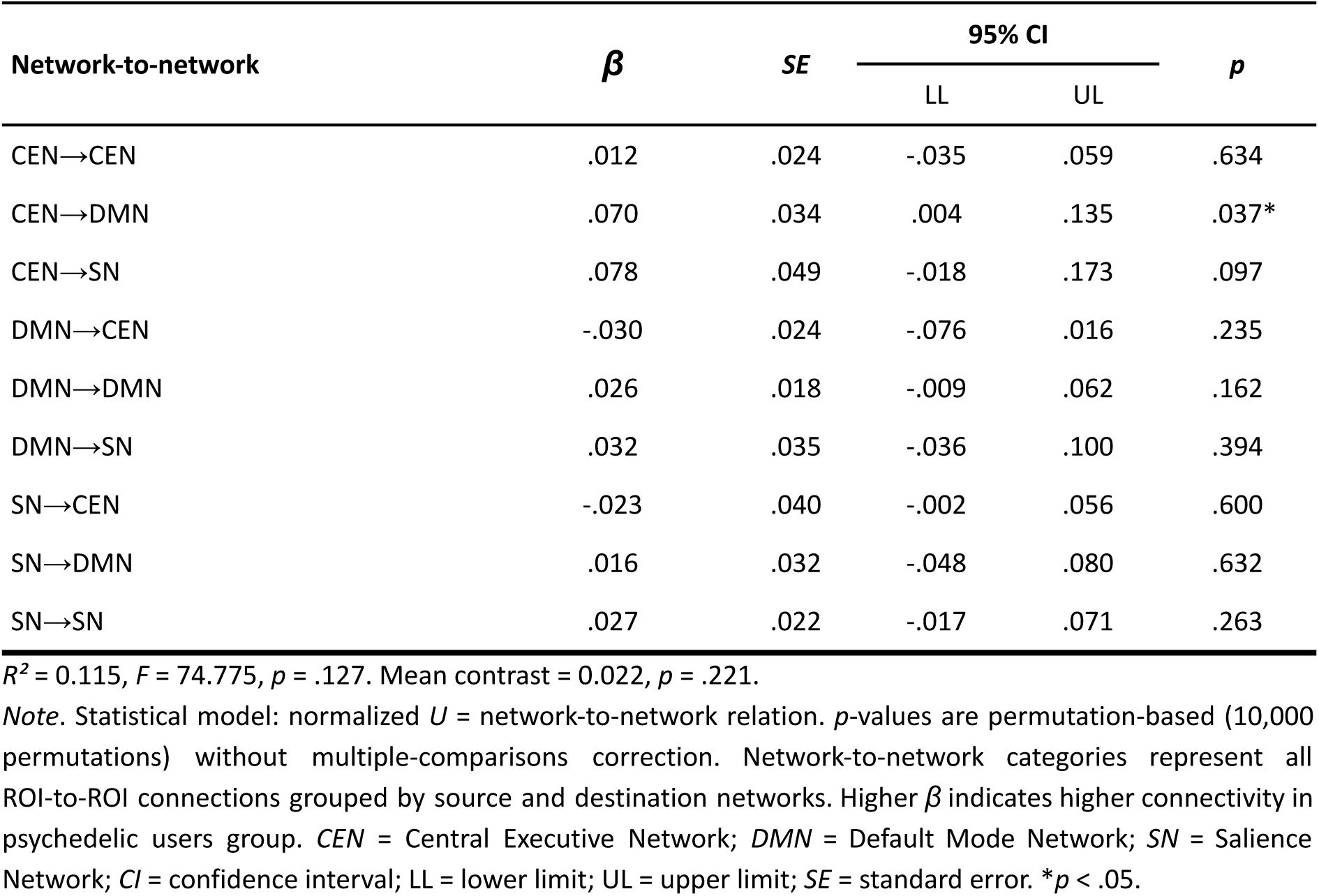
Regression Analysis Predicting Normalized U in Network-to-Network Directed Connections.

| Network-to-network | $\beta$ | SE | 95% CI | | $p$ |
| --- | --- | --- | --- | --- | --- |
|  |  |  | LL | UL |  |
| CEN→CEN | .012 | .024 | -.035 | .059 | .634 |
| CEN→DMN | .070 | .034 | .004 | .135 | .037* |
| CEN→SN | .078 | .049 | -.018 | .173 | .097 |
| DMN→CEN | -.030 | .024 | -.076 | .016 | .235 |
| DMN→DMN | .026 | .018 | -.009 | .062 | .162 |
| DMN→SN | .032 | .035 | -.036 | .100 | .394 |
| SN→CEN | -.023 | .040 | -.002 | .056 | .600 |
| SN→DMN | .016 | .032 | -.048 | .080 | .632 |
| SN→SN | .027 | .022 | -.017 | .071 | .263 |

$R^2 = 0.115$ , $F = 74.775$ , $p = .127$ . Mean contrast = 0.022, $p = .221$ .
*Note.* Statistical model: normalized $U$ = network-to-network relation. $p$ -values are permutation-based (10,000 permutations) without multiple-comparisons correction. Network-to-network categories represent all ROI-to-ROI connections grouped by source and destination networks. Higher $\beta$ indicates higher connectivity in psychedelic users group. CEN = Central Executive Network; DMN = Default Mode Network; SN = Salience Network; CI = confidence interval; LL = lower limit; UL = upper limit; SE = standard error. \* $p < .05$ .

Only one connection reached statistical significance, though with a small effect size and falling outside our preregistered hypotheses: CEN-to-DMN (*β* = .070, *p* = .037). We additionally conducted explanatory interaction analyses (Model 2: contrast ∼ network_relation × bands; Models 3 and 4: contrast ∼ city + eyes + network_relation × bands; with and without dataset merging), three of which reached nominal statistical significance (see Supplementary Material 4, Table S4). However, none of the described effects survived correction for multiple comparisons.

Moreover, to visualize between-group connectivity comparisons across all ROIs and conditions, we conducted an analysis inspired by specification curve methodology. Figure 4 displays standardized effect sizes for all 9,240 unique comparisons, each representing a distinct combination of condition, frequency band, network-to-network relation, and dataset. The specification curve remained remarkably flat (*M* = 0.023, *SD* = 0.022), with no visible clusters across specifications and no outliers representing connectivity changes beyond those predicted by our predefined networks. This demonstrates that our null results are robust across the entire range of ROIs, rather than being obscured by our network grouping approach.

**Figure 4.**
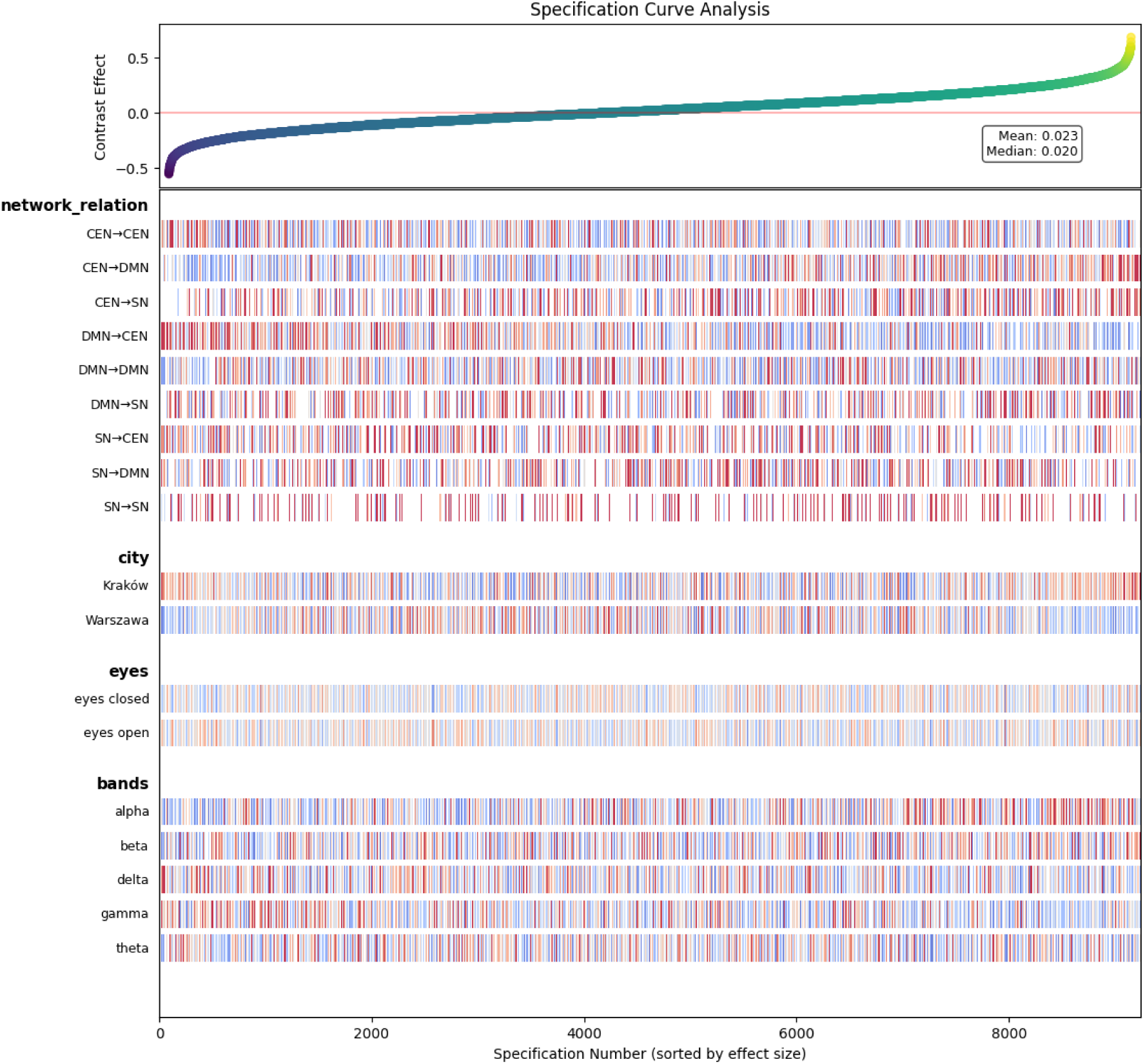
Specification Curve Analysis. Upper panel: Standardized effect sizes (Mann-Whitney *U*) for 9,240 connectivity comparisons ordered by magnitude (shaded area = 95% CI). Lower panels: analytical specifications. Higher *U* values indicate higher connectivity in psychedelic users. *CEN* = Central Executive Network; *DMN* = Default Mode Network; *SN* = Salience Network; Kraków = dataset I; Warszawa = dataset II.

## Discussion

We examined whether long-term naturalistic psychedelic users exhibit distinctive resting-state brain activity patterns compared to matched non-users. This larger-scale cross-sectional investigation of individuals reporting repeated psychedelic use who remained abstinent for at least 30 days prior to data collection contributes a crucial perspective for addressing current knowledge gaps in this field (Aday et al., 2020; Carhart-Harris, 2018; Gattuso et al., 2023).

We analysed oscillatory power, signal complexity, and effective connectivity using complementary analytical methods and observed predominantly null results. Although condition-specific differences emerged, effect sizes were small and mostly inconsistent — both internally (showing no coherent pattern) and relative to our a priori hypotheses. Below, we discuss how these findings relate to acute psychedelic effects, long-term repeated exposure in naturalistic users, and existing knowledge gaps.

### Main findings

Oscillatory power analysis revealed no significant main effects of the group in any frequency band, indicating comparable overall power between groups. However, significant group × condition interactions emerged (in theta, beta, and gamma). Psychedelic users exhibited lower PSD values in gamma compared to non-users in eyes-open condition. Signal complexity analyses showed no significant main effect of the group. However, psychedelic users exhibited significantly lower LZ scores compared to non-users in eyes-open condition (*t* = 2.33, *p* = .022). Effective connectivity analyses revealed no significant group differences in network connectivity across frequency bands. Although isolated effects of small magnitude were observed (CEN-to-DMN and DMN-to-CEN × gamma interaction), none survived corrections for multiple comparisons. Our findings did not show any hypothesized neurophysiological signatures, derived from reports of several acute and post-acute clinical trials, chronic ayahuasca users and selected mechanistic theories of psychedelic action.

On the other hand, our results align to some extent with those observed in one clinical trial involving single-dose psilocybin administration by McCulloch et al. (2022). They demonstrated minimal persistence of substance-related neurophysiological effects, particularly at longer follow-up intervals. More specifically, healthy psychedelic-naïve volunteers exhibited significant reductions in Executive Control Network connectivity at 1 week, but these effects had dissipated by 3 months. Notably, every other within- and between-network connectivity measures showed small to negligible effect sizes at both follow-up timepoints. Despite a robust methodological approach, their trial’s inferential capacity remains limited by the small sample size (n=10) .

### Mechanistic models

The mechanistic frameworks evaluated here — the EBH, REBUS, and neuroplasticity models — currently have substantial empirical support from acute, controlled settings. A critical distinction must therefore be drawn: the studies underlying these models predominantly examine rigorous psychedelic administration protocols (fixed doses, pharmaceutical-grade compounds, structured preparation and integration sessions), whereas our study examines uncontrolled naturalistic use patterns. Rather than directly testing these models, our aim was to explore whether their proposed neurophysiological mechanisms extend to a fundamentally different context: repeated self-administration over extended time periods, with current abstinence during study sessions.

The EBH posits that psychedelics acutely increase brain complexity, signal diversity, and criticality (Carhart-Harris, 2018; Carhart-Harris et al., 2014); we observed reduced signal complexity in long-term naturalistic psychedelic users. The REBUS model, which emerged from the EBH, posits the possibility of persistent changes in neural excitability and higher-order network dynamics, particularly reduced DMN integrity (Carhart-Harris & Friston, 2019); our analyses revealed no differences in within- or between-network connectivity patterns. The TNT posits that several psychopathologies stem from disrupted DMN-SN-CEN balance (B. Menon, 2019; V. Menon, 2011; V. Menon & Uddin, 2010), and altered interactions among these networks have been implicated as neurophysiological markers predicting clinical outcomes (Whitfield-Gabrieli & Ford, 2012) and informing transdiagnostic frameworks (Insel et al., 2010). Psychedelic interventions reportedly alter DMN-SN-CEN connectivity — reducing within-DMN connectivity, enhancing SN-to-DMN connectivity, or producing broader desynchronization (Gattuso et al., 2023; Knudsen, 2023; Madsen et al., 2021; Siegel et al., 2024; Stoliker et al., 2023; Thomas et al., 2017); we found no supporting evidence of such network-level changes in long-term psychedelic users. Neuroplasticity-based models suggest that psychedelics induce lasting effects through sustained reorganization of neural circuit dynamics (Calder & Hasler, 2023; Dos Santos et al., 2016; Grieco et al., 2022; Ly et al., 2018; Siegel et al., 2024; Vollenweider & Kometer, 2010); apart from complexity, our sample did not exhibit significant differences in neural dynamics as detectable within the sensitivity of the methods employed.

The absence of group differences does not invalidate these mechanistic models but rather contributes novel constraints on their scope, highlighting potential boundary conditions of psychedelic effects. A crucial question emerges: can results from controlled, limited-dose clinical trials and inferences derived from acute and post-acute states be translated into the neurobiological signatures of repeated naturalistic use? Our study cannot provide definitive answers but rather underscores the need to consider whether these changes are specific to acute and short-term post-acute states, to clinical populations, or whether they reflect the sensitivity of particular methodological choices and measures.

### Exploratory or unexpected results

Although our hypotheses were mostly not confirmed, we observed some unanticipated results. Psychedelic users exhibited lower signal complexity during the eyes-open condition, and exploratory analyses of oscillatory power revealed altered magnitudes in eyes-open versus eyes-closed contrasts. Several interpretive frameworks may be considered.

The condition-specificity of this effect — present in eyes-open but absent in eyes-closed — can be itself informative. It can, for example, suggest the group difference is not attributable to a global trait-level shift in neural dynamics but rather to state-dependent processing differences emerging during active visual input. This observation resonates with findings from Mediano et al. (2024), which demonstrated that environmental conditions dramatically modulate psychedelic-related entropy changes: LSD-induced complexity increases were largest with eyes closed and substantially diminished under external stimulation (eyes-open, music, or video). Our group difference thus emerged precisely under the conditions where LZ complexity appears most sensitive to contextual modulation. One possible interpretation is that long-term psychedelic users process external visual input in a more constrained or less variable manner during the eyes-open resting state. Whether this reflects lasting neurobiological changes — such as subtle, long-term consequences of repeated 5-HT2A receptor activation on cortical processing (discussed below) — or other factors such as differences in attentional habits, visual processing strategies, or pre-existing traits associated with psychedelic use, cannot be determined from the present data.

Alternative explanations including methodological artifacts, measurement noise, or Type I error cannot be excluded. Given the small effect size and the absence of convergent findings from other measures in the present study, we refrain from strong mechanistic interpretations. Hypothesis-driven studies designed to examine condition-dependent complexity patterns and their relationship to psychedelic use history should determine whether this effect is replicable and functionally meaningful.

### Distinguishing acute and long-term effects

5-HT2A receptor activation is the primary molecular mechanism mediating psychedelic effects (Madsen et al., 2019; Nichols, 2016; Preller et al., 2018). The 5-HT2A receptor plays a central role in modulating cortical excitability, network dynamics, and functional connectivity particularly within the DMN (Beliveau et al., 2017; Gattuso et al., 2023; Knudsen, 2023). Critically, repeated agonism of this receptor leads to its downregulation: in animal models, repeated administration of psychedelics produces diminished 5-HT2A receptor density in frontal cortex (Buckholtz et al., 1988; De La Fuente Revenga et al., 2022), and in humans, chronic ayahuasca users show significantly reduced neocortical 5-HT2A receptor binding compared to controls (Dos Santos et al., 2016). This downregulation is thought to underlie psychedelic tolerance — a phenomenon whereby subjective effects diminish rapidly with repeated administration (Nichols, 2016). Yet the downstream consequences of such receptor adaptation for neural dynamics remain poorly understood, particularly in the context of long-term naturalistic use (Grieco et al., 2022; Wallach et al., 2023).

While acute psychedelic administration promotes neuroplasticity through 5-HT2A activation (Calder & Hasler, 2023; Ly et al., 2018) chronic exposure may reduce neuroplastic capacity through compensatory downregulation — potentially attenuating the neurobiological substrate for psychedelic-induced effects. Reduced 5-HT2A-mediated modulation of cortical dynamics could decrease baseline signal complexity, consistent with the lower LZ values we observed in the eyes-open condition. Our predominantly null findings in connectivity may reflect a return to the brain’s baseline state after acute effects resolve, even following repeated use. Receptor downregulation, homeostatic reequilibration, or simply the passage of sufficient time for all post-acute effects to dissipate could each contribute to explaining both our findings and those from single-administration studies reporting minimal persistence of neurophysiological effects (McCulloch et al., 2022) — though such accounts remain speculative and need to be confirmed in future studies. The extent to which chronic users and single-dose clinical trial participants differ neurobiologically — in circuit-level organization, receptor signaling, neuroendocrine function, or genetics — remains largely unexplored, yet could itself inform models positing lasting neural reorganization (Dos Santos et al., 2016; Grieco et al., 2022; Knudsen, 2023).

### Methodological considerations

Our findings should also be interpreted within the broader context of methodological challenges in psychedelic research. Van Elk and Fried (2023) concluded that most psychedelic clinical trials “lack basic quality controls and methodological rigor,” identifying pervasive validity threats including inadequate sample sizes, analytical flexibility, publication bias or failed blinding (participants identifying allocation with >90% accuracy). Neuroimaging studies often rely on small samples (10-20 participants) including datasets re-analyzed across multiple publications (as McCulloch et al., 2022 reported: 52% of published literature on psychedelic effects assessed with resting-state fMRI was based on two datasets), threatening external and statistical validity of those reports (Gattuso et al., 2023; van Elk & Fried, 2023). Moreover, null results could be in fact substantially underreported, hence creating inflated impressions of robust effects.

Null findings have been consistent across nearly all analyses of the present dataset, despite employing different analytical approaches (Ruban et al., 2025). One exception emerged: event-related potential analyses revealed group differences in early emotional processing (N170, N200) but not in the higher-order cognitive components (P200, P300; Orłowski et al., 2024). We suggest this may be reflecting superior signal-to-noise ratio in task-based versus resting-state paradigms. Nevertheless, this absence of group differences in higher-order cognitive components is consistent with our resting-state connectivity findings, given that the DMN, CEN, and SN are among the most extensively studied neural substrates of higher-order cognition and conscious processes.

In light of these field-wide concerns, the Siegel et al. (2024) findings — among the most prominent recent reports of lasting psychedelic effects — also warrant methodological consideration. Notably, their sample comprised 7 healthy volunteers (one of whom was a co-author of the study), the first author served as a facilitator tasked with building therapeutic alliance, and the replication sample was reduced to 4 participants due to attrition. Such observations, rather than invalidating individual studies, illustrate the broader need for independent replication with adequately powered, pre-registered designs.

Present study represents a larger-scale examination than is typical in psychedelic neuroimaging research. Small samples combined with inadequately addressed multiple comparisons substantially undermine statistical power (McCulloch et al., 2022; van Elk & Fried, 2023). To maximize inferential capacity within our design, we attempted formal a priori sample size estimation and post-hoc power analysis. However, these attempts proved unviable due to insufficient reporting of effect sizes in the existing literature and the limited availability of comparable research, which precluded reliable estimation. Therefore, despite recruiting a larger sample than is typical in this literature, we cannot determine whether we achieved adequate statistical power. More subtle neurophysiological differences, particularly those underlying higher-order cognition, may remain undetectable above population-level noise without substantially larger cohorts or longitudinal designs.

Participant selection and baseline neural state represent additional considerations. Clinical trials recruiting participants with psychiatric diagnoses may effectively study brain dynamics with atypical baseline connectivity — for example, DMN hyperconnectivity in depression (Whitfield-Gabrieli & Ford, 2012). Consequently, the prominent connectivity changes observed in such trials could reflect normalization of pathological patterns rather than alterations to healthy neural architecture. Furthermore, clinical trials are susceptible to expectancy effects, ineffective placebo control, derandomization, and attrition bias — factors that can substantially influence observed effect sizes and inferential capacity (van Elk & Fried, 2023).

The well-established context-dependency of psychedelic effects raises further questions. Specific laboratory conditions — such as lying still in a confined MRI scanner, either blindfolded or with eyes open — could potentially amplify or otherwise modulate certain neurobiological signatures. This possibility, though long recognized (e.g. Strassman, 1991). remains rarely discussed. Concerns have also been raised about the interpretation and comparability of neurophysiological effects obtained under different experimental paradigms, with even the simple distinction between eyes-open and eyes-closed conditions frequently overlooked (McCulloch, Knudsen, et al., 2022)

For this reason, we employed both eyes-closed and eyes-open conditions, whereas most studies employ only one (Agcaoglu et al., 2019; McCulloch, Knudsen, et al., 2022). Our results demonstrate evidence that oscillatory power, complexity, and effective connectivity patterns vary significantly between these conditions. Observed consistency of null findings across both conditions in our between-group comparisons strengthens our confidence that the absence of effects is in this case not attributable to condition-specific factors.

Finally, the high diversity of contexts characterizing naturalistic use — varying set and setting, dosing regimens, substance purity, concurrent practices such as meditation or integration — remains largely unaddressed in neuroimaging research, though these factors could serve as boundary conditions for testing the moderators of psychedelic-related effects (Gonzalez-Mulé & Aguinis, 2018; Hartogsohn, 2016). Mediano et al., (2024) demonstrated that environmental conditions, including music and video, dramatically reduce the magnitude of LSD-induced changes in neural complexity and desynchronization compared to eyes-closed resting-state conditions. Our findings are consistent with context-dependency, although the present study’s cross-sectional design precludes disentangling the influence of potentially confounding factors or establishing causal relationships.

### Future directions

We conclude by echoing existing calls for methodological pluralism in psychedelic research. Carhart-Harris et al. (2022) stressed the need for “pragmatic research” — including digital phenotyping, remote monitoring technologies, and large-scale naturalistic studies — as complementary evidence streams alongside confirmatory clinical trials. Similarly, Schenberg (2024) argued that real-world evidence and diverse methodological approaches should be incorporated into decision-making and regulatory processes, contending that conventional procedures may be too narrow to adequately capture the complexity of psychedelic-assisted interventions. Maintaining a critical stance toward simplified mechanistic theories and trial designs, while embracing multifactorial models supported by inclusive participant samples and diverse methodologies, seems essential for advancing an understanding of psychedelic effects that is both scientifically rigorous and clinically or socially relevant.

### Limitations

Our study is not without several limitations. Cannabis use, though largely matched between groups, could not be fully assessed due to incomplete data collection, representing a significant confound given cannabis effects on brain activity. Meditation practice, which can produce similar DMN changes as psychedelics (Smigielski et al., 2019), may have obscured group differences. Naturalistic psychedelic use occurs within highly variable set and setting conditions, including: dosage variability, substance purity, co-administered substances, social context, environmental conditions, and psychological preparation differ across experiences and may influence neurobiological outcomes in ways we are not aware of. Furthermore, our self-selected and fully Polish sample may not represent the broader population of psychedelic users.

Technical limitations include EEG’s inherent spatial resolution constraints, reliance on standardized rather than individual brain templates, and limited sensitivity to subcortical structures potentially relevant to psychedelic action. Our LZ analysis employed four-second epochs, consistent with common practice but without established guidelines for optimal window length. LZ complexity analysis might therefore not capture network-level or cross-channel dynamics relevant to large-scale brain organization. The interpretation of complexity values remains challenging, as elevated complexity may reflect either richer information processing or greater signal noise. In oscillatory power, significant site differences emerged in theta, beta, and gamma bands, likely reflecting differences in recording equipment, electrode configurations, sampling rates, or participant characteristics between datasets. Although preprocessing addressed these differences, residual effects may persist. Moreover, the reliance on Fourier-based spectral analysis assumes signal stationarity and sinusoidal decomposition, which does not capture rapid, transient brain activity. Our connectivity analysis relied on 22 a priori ROIs that may have not fully captured critical nodes. Also, using nDTF has inherent limitations including sensitivity to preprocessing choices, epoch length, and frequency band definitions. Our source-localisation connectivity analysis relied on a standardized brain template without participant-specific structural scans, introducing variability in source reconstruction accuracy across individuals. Our source-localization of EEG signals also precluded analysis of subcortical structures including the thalamus, claustrum, amygdala, and hippocampus, which play important roles in psychedelic action. The conventional frequency band definitions may not optimally capture individual differences in peak frequencies or psychedelic-specific signatures. The TNT, however influential, represents a simplified model and as, for example, Gattuso et al. (2023) noted: questions remain about what DMN modulation represents mechanistically (while conceptual flexibility allows broad application of network models, it also risks overinterpretation unless supported by more precise operational definitions and mechanistic links). Lastly, our cross-sectional design cannot rule out the possibility that observed differences reflect pre-existing group characteristics related to psychedelic use rather than consequences of the substances themselves, nor can it determine whether effects of psychedelics emerge or dissipate over different timescales. Other whole-brain connectivity analyses, and using alternative methods on these datasets also yielded mostly null results (Orłowski et al., 2024a; Ruban et al., 2025). Nevertheless, null effects in effective connectivity may partly reflect limited signal-to-noise ratio and spatial precision inherent to EEG-based source connectivity estimates.

### Closing summary

Our large-sample investigation of naturalistic psychedelic users revealed predominantly null findings across oscillatory power, signal complexity, and effective connectivity. These results highlight three considerations: (1) long-term effects of limited exposure must be distinguished from chronic repeated use, which may involve opposing receptor dynamics and distinct neurobiological adaptations; (2) context-dependency and baseline neural states substantially influence the expression of psychedelic effects; (3) mechanistic frameworks should accommodate homeostatic compensation and population heterogeneity. While not confirming predictions derived from current theoretical models, our findings provide valuable constraints for developing a more nuanced understanding of lasting psychedelic neurobiological mechanisms. Importantly, these results do not directly challenge existing models of acute psychedelic action, but rather indicate that such models may not straightforwardly generalize to long-term, post-acute, naturalistic use contexts.

## Data Availability

All data and code generated during this study are openly available. Raw behavioral data, EEG data, source code, analysis scripts (Python, MATLAB, R) with proper documentation are provided in a zenodo repository (https://doi.org/10.5281/zenodo.16534101). The study was preregistered at OSF (https://osf.io/z6tky). Every contact on replication, re-analysis, methods extensions or any other thoughts regarding this project is welcome.

## Acknowledgements

We thank Anastasiia Ruban (SWPS University) for her broad contributions making this whole project possible. Her commitment towards inclusive and mindful psychedelic science has been an inspiration throughout this work. We also thank Mirosław Wyczesany (Jagiellonian University) for sharing the ASCT with us, while it was still under development, and for all the time he dedicated to consult technicals of our nDTF analyses.

## Authors Contribution (CRediT)

Conceptualization: MW, MWi, PL, PO

Data curation: PL, PO

Formal analysis: MW, PL, PO, SA

Funding acquisition: JH, MWi

Investigation: PO, SA, JH

Methodology: MW, PL, PO

Project administration: JH

Resources: MWi

Software: PL

Supervision: MB, MWi

Validation: JH, PL, PO

Visualization: MW, PL, PO, SA

Writing – original draft: MW (lead), PL, PO, SA

Writing – review & editing: MW, MWi, MB, PL, PO, SA, JH

