## Supplementary Materials for "When the psychedelic state’s over: limited evidence for persistent neurophysiological changes in naturalistic psychedelic users"

### Supplementary Material 1

**Table S1.** *Participant Demographics, Substance Use, and Meditation Practice by Dataset and Group.*

|  | **Dataset I (Kraków)** | |  | **Dataset II (Warszawa)** | |  |
| --- | --- | --- | --- | --- | --- | --- |
|  | **Psychedelic users  (n=21)** | **Non-users  (n=19)** | **Between-group comparisons** | **Psychedelic users^1)^  (n=32)** | **Non-users  (n=30)** | **Between-group comparisons** |
| **Demographics** |  |  |  |  |  |  |
| Age  (years) | 26 (6) | 29 (7) | t(33)= -1.31  n.s. | 29 (5) | 28 (4) | t(59)= -0.61 n.s. |
| Sex (male/female) | 12 / 9 | 10 / 9 | χ²(1) = 0.08  n.s. | 9 / 23 | 10 / 20 | χ²(1) = 0.20  n.s. |
| **Education Level**  (finished) |  |  | χ²(3) = 0.41  n.s. |  |  | χ²(3) = 2.54  n.s. |
| Secondary | 10 (48%) | 8 (42%) |  | 12 (33%) | 7(23%) |  |
| Bachelor’s | 3 (14%) | 3 (16%) |  | 8 (22%) | 8 (27%) |  |
| Master’s | 7 (33%) | 6 (32%) |  | 11 (31%) | 15 (50%) |  |
| Above  Master’s | 1 (5%) | 2 (10%) |  | 1 (3%) | 0 (0%) |  |
| **Substance Use** |  |  |  |  |  |  |
| Lifetime psychedelics | 38 (38) | — | — | 26 (11) | — | — |
| Lifetime cannabis^2)^ | 160 (305) | 267 (588) | t(14) = 0.54  n.s. | 324 (316) | 92 (128) | t(25) = 3.17** |
| Alcohol  (AUDIT score) | 2 (1) | 3 (2) | t(36) = –1.26 n.s. | 3 (2) | 3 (2) | t(57) = –0.83  n.s. |
| **Meditation** |  |  |  |  |  |  |
| Lifetime practice  (hours) | 763 (2183) | 246 (794) | t(26) = 1.01  n.s. | 381 (920) | 136 (371) | t(41) = 1.39  n.s. |
| *Note. Data are presented as M (SD) for continuous variables and n (%) for categorical variables. Statistical comparisons performed using independent samples Welch's t-test for continuous variables and χ² test for categorical variables (two-tailed, α = .05). AUDIT* = Alcohol Use Disorder Identification Test.  *ªn* = 4 missing data. *ᵇ* Lifetime cannabis data incomplete due to programming error (in dataset I: n = 13 users, n = 11 non-users; dataset II: n = 20 users, n = 26 non-users cannabis data). *n.s.* = non-significant. ***p* < .01. | | | | | | |

#### Laboratory session details

Each EEG session began with documentation review, verbal instructions, and written informed consent procedures, followed by cap preparation for EEG (approximately 45 min total). Participants then completed three short questionnaires: Reflection-Rumination Questionnaire (RRQ) State-Trait Anxiety Inventory (STAI) and Beck Depression Inventory-II (BDI-II).

Subsequently, participants were seated comfortably in front of a computer screen (approximately 60 cm’s) and instructed to: (1) maintain their gaze on a central fixation cross for 5 min (eyes-open condition), then (2) close their eyes and relax for 5 min (eyes-closed condition). Signals from both earlobes were recorded. Four electrooculograms (EOG) were recorded using bipolar electrodes: two vertical (VEOG), above and below the left eye (supra- and infra-orbital) and two horizontal (HEOG), at the external canthi of both eyes. After the resting-state recording, participants completed the Amsterdam Resting-State Questionnaire (ARSQ) to assess e.g. contents of their thoughts/subjective experiences during recording.

EEG sessions concluded with three additional experimental paradigms (for more details of these tasks see preregistration at <https://osf.io/z6tky>; Hobot et al., 2021; or published analyses at Orłowski et al., 2024b, 2024a; Ruban et al., 2025). Participants were frequently offered short breaks between procedure steps/within tasks and always provided as needed. Total session duration was approximately 2.5 hr. None of the participants withdrew during laboratory sessions and none of them declared experiencing unwellness/undesired symptoms or any other negative reactions.

#### Questionnaire and Substance Use Data Visualization

Split violin plots display score distributions between groups. Violin shapes represent probability density distributions, with wider sections indicating higher data density. Overlaid boxplots show median (central line), interquartile range (box), and whiskers extending to 1.5×IQR. Individual data points displayed as jittered dots. Statistical comparisons performed via independent samples t-test (two-tailed, α = .05). P-values less than .001 may be reported as *p* < .001 rather than exact values.

##### Dataset I (Kraków)

| 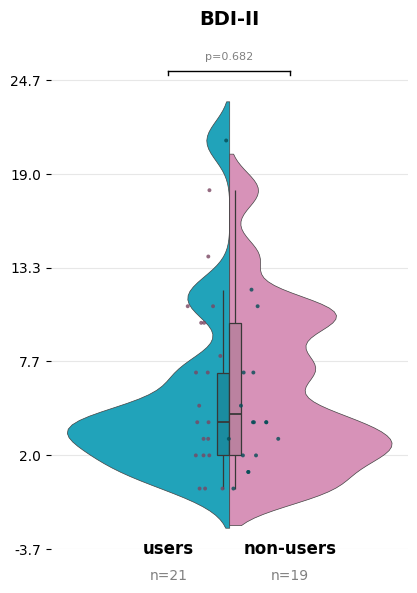 | 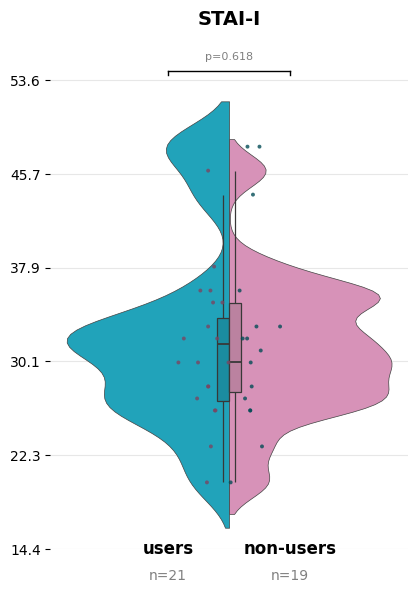 | 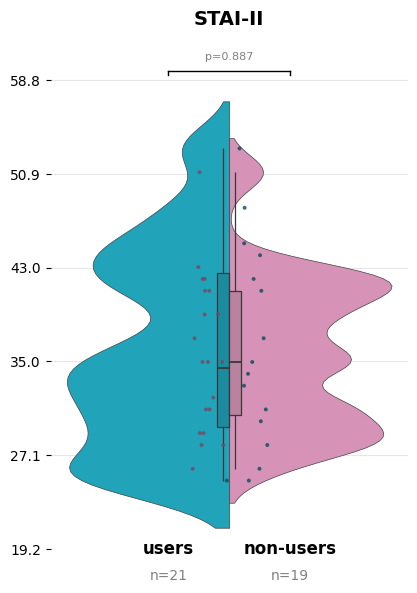 |
| --- | --- | --- |

###### Figure S1. *State-Trait Anxiety and Depression Inventory Score Distributions by Group (Dataset I).* Violin shapes represent probability density distributions; boxplots show median (center line), interquartile range (box), and whiskers (1.5× IQR). Individual data points shown as jittered dots (psychedelic users = blue/left; non-users = pink/right). Error bars represent range between first and third quartiles. STAI-I = State Anxiety Inventory; STAI-II = Trait Anxiety Inventory; BDI-II = Beck Depression Inventory-II; *n* = sample size for each group.

| 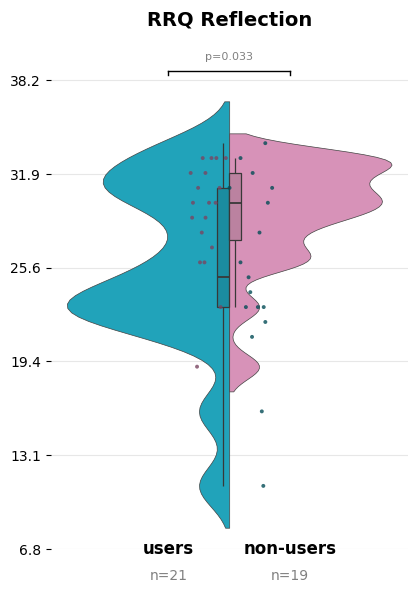 | 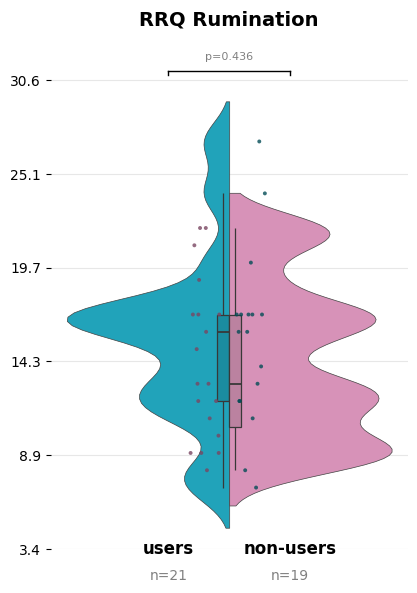 |
| --- | --- |

###### Figure S2. *Reflection-Rumination Questionnaire Subscale Distributions by Group (Dataset I).* Violin shapes represent probability density distributions; boxplots show median (center line), interquartile range (box), and whiskers (1.5× IQR). Individual data points shown as jittered dots (psychedelic users = blue/left; non-users = pink/right). Error bars represent range between first and third quartiles. RRQ = Reflection-Rumination Questionnaire; *n* = sample size for each group.

| 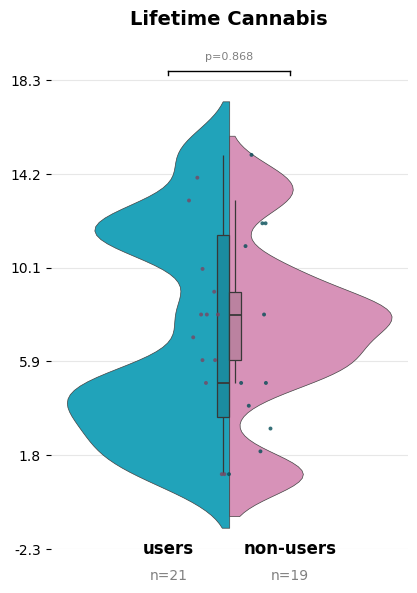 | 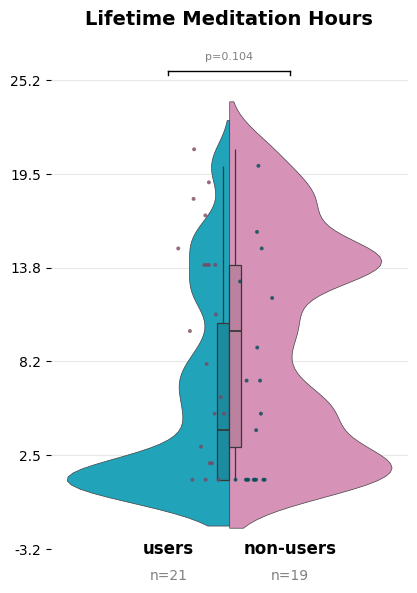 |
| --- | --- |

**Figure S3.** *Ranked Lifetime Cannabis Use and Meditation Hours Distributions by Group (Dataset I).* Raw frequency data were converted to dense ranks to reduce the influence of extreme outliers and create ordinal scales suitable for group comparisons. Violin shapes represent probability density distributions of ranks for each group, with wider sections indicating higher data density at those rank levels. Boxplots show median (center line), interquartile range (box), and whiskers (1.5× IQR). Individual data points shown as jittered dots (psychedelic users = blue/left; non-users = pink/right). *n* = sample size for each group.

| 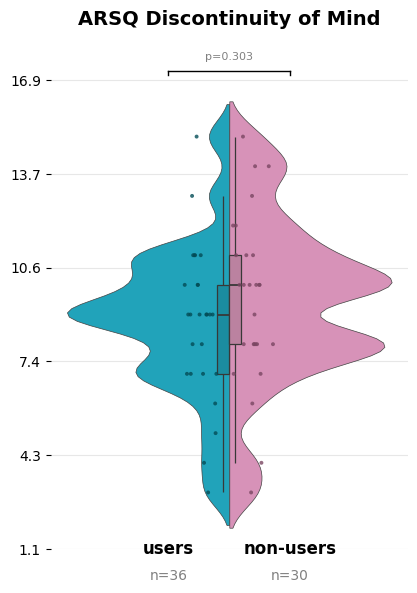 | 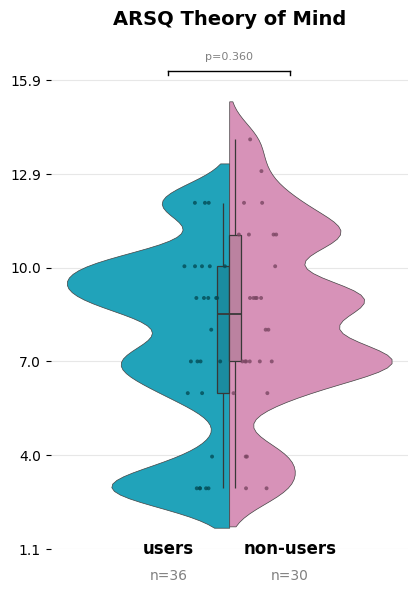 | 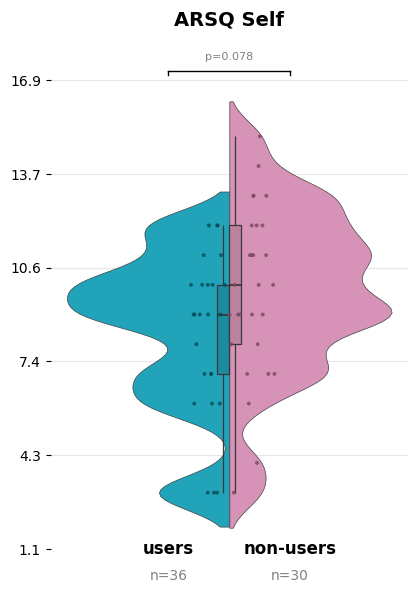 | 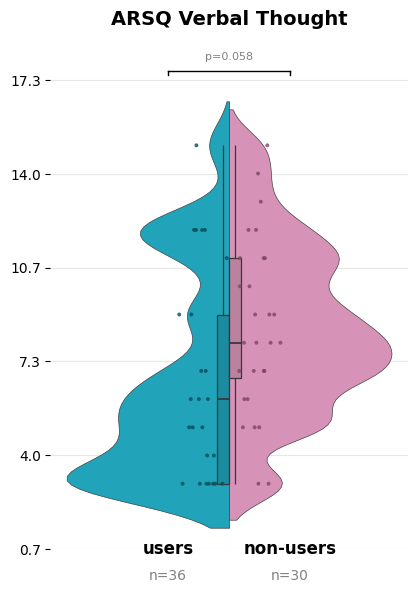 |
| --- | --- | --- | --- |
| 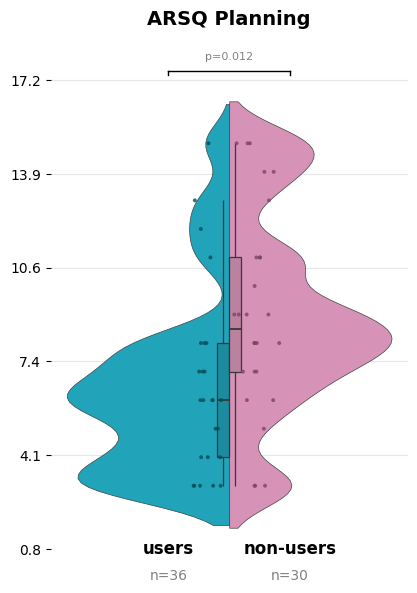 | 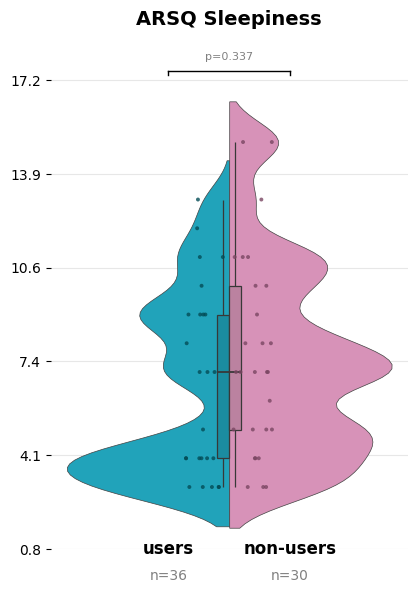 | 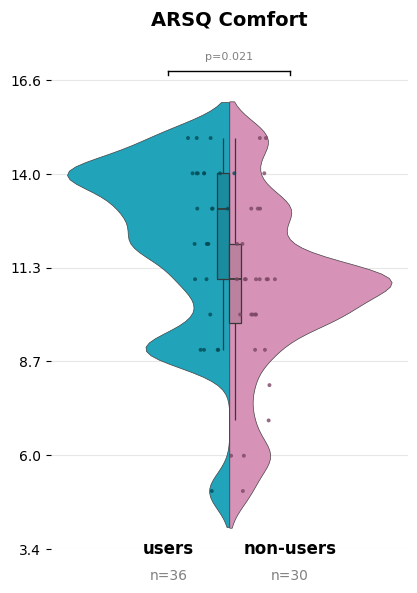 | 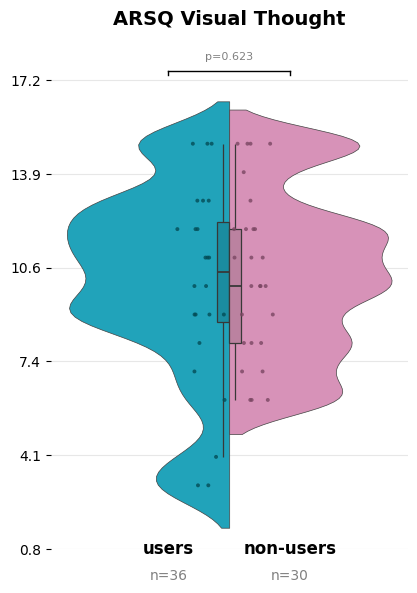 |
| 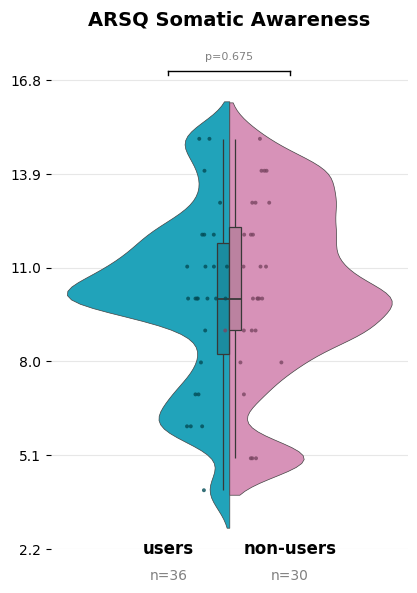 | 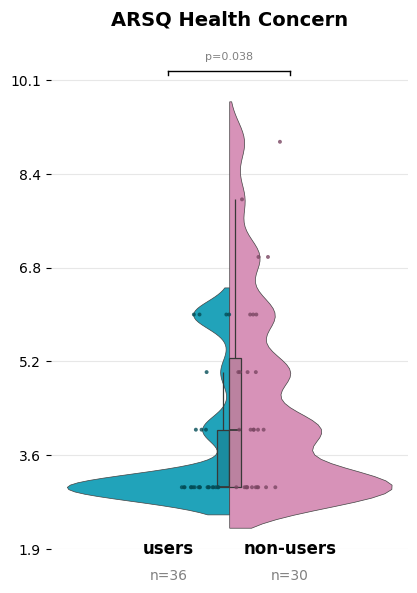 |  |  |

**Figure S4.** *Amsterdam Resting-State Questionnaire Subscale Distributions by Group (Dataset I).* Violin shapes represent probability density distributions; boxplots show median (center line), interquartile range (box), and whiskers (1.5× IQR). Individual data points shown as jittered dots (psychedelic users = blue/left; non-users = pink/right). Error bars represent range between first and third quartiles. ARSQ = Amsterdam Resting-State Questionnaire; *n* = sample size for each group.

##### Dataset II (Warszawa)

| 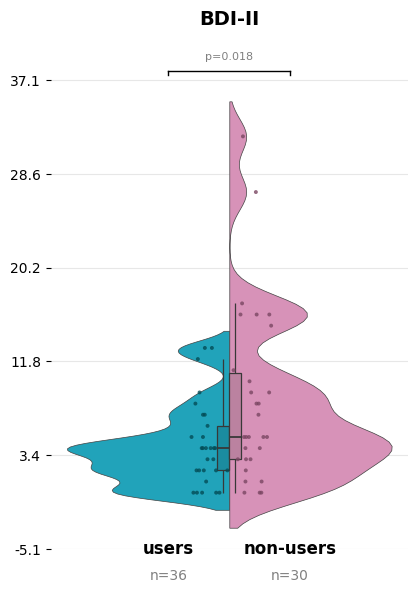 | 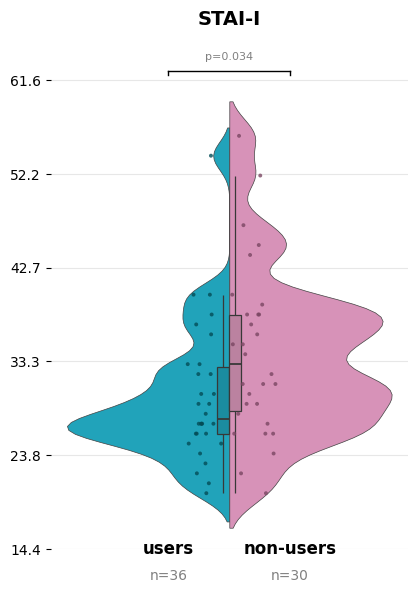 | 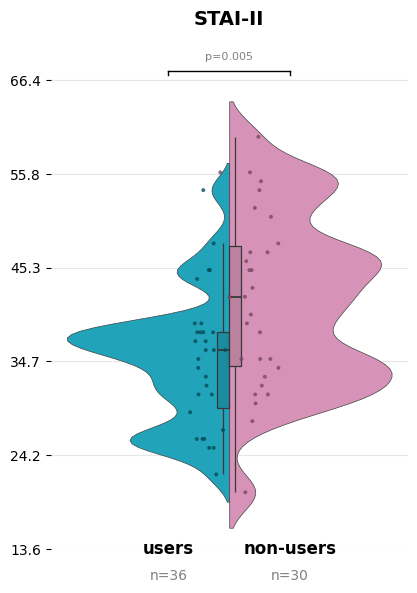 |
| --- | --- | --- |

**Figure S5.** *State-Trait Anxiety and Depression Inventory Score Distributions by Group (Dataset II).* Violin shapes represent probability density distributions; boxplots show median (center line), interquartile range (box), and whiskers (1.5× IQR). Individual data points shown as jittered dots (psychedelic users = blue/left; non-users = pink/right). Error bars represent range between first and third quartiles. STAI-I = State Anxiety Inventory; STAI-II = Trait Anxiety Inventory; BDI-II = Beck Depression Inventory-II; *n* = sample size for each group.

| 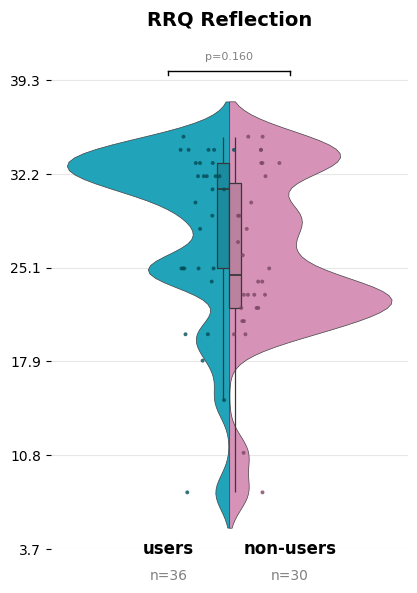 | 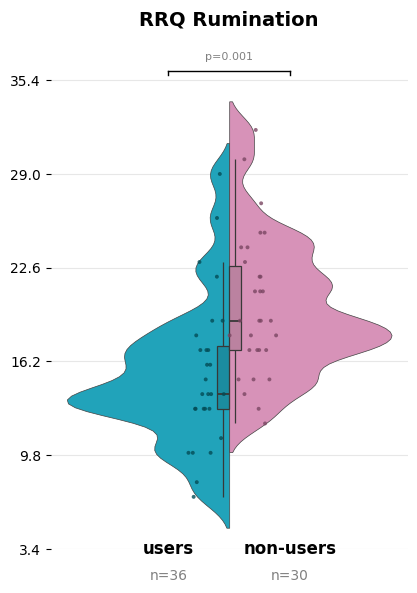 |
| --- | --- |

###### Figure S6. *Reflection-Rumination Questionnaire Subscale Distributions by Group (Dataset II).* Violin shapes represent probability density distributions; boxplots show median (center line), interquartile range (box), and whiskers (1.5× IQR). Individual data points shown as jittered dots (psychedelic users = blue/left; non-users = pink/right). Error bars represent range between first and third quartiles. RRQ = Reflection-Rumination Questionnaire; *n* = sample size for each group.

| 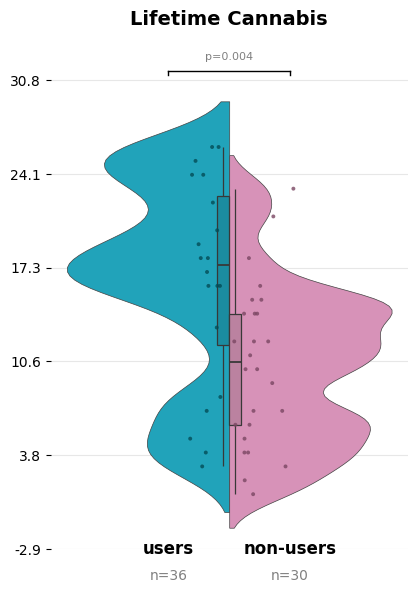 | 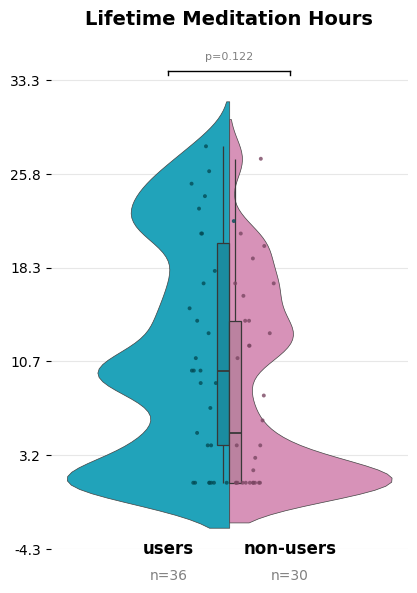 |
| --- | --- |

**Figure S7.** *Ranked Lifetime Cannabis Use and Meditation Hours Distributions by Group (Dataset II).* Raw frequency data were converted to dense ranks to reduce the influence of extreme outliers and create ordinal scales suitable for group comparisons. Violin shapes represent probability density distributions of ranks for each group, with wider sections indicating higher data density at those rank levels. Boxplots show median (center line), interquartile range (box), and whiskers (1.5× IQR). Individual data points shown as jittered dots (psychedelic users = blue/left; non-users = pink/right). *n* = sample size for each group.

| 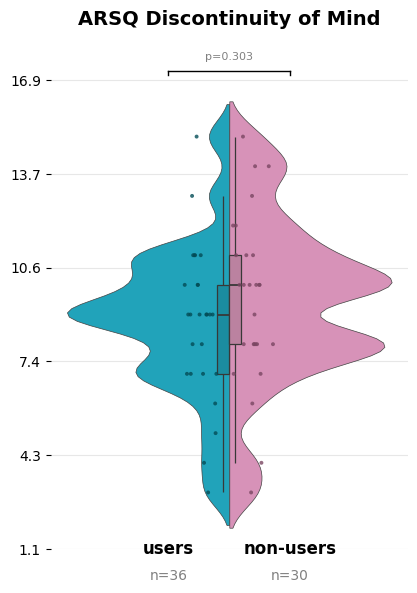 | 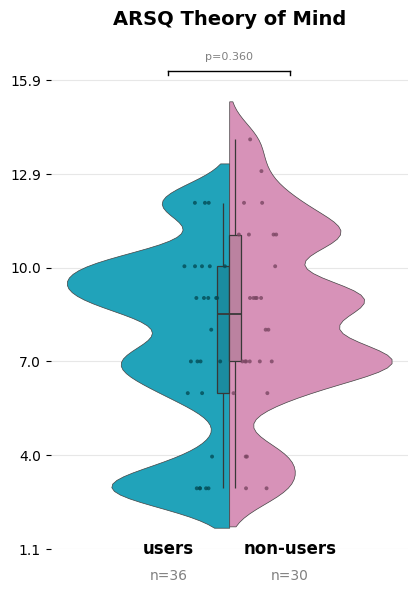 | 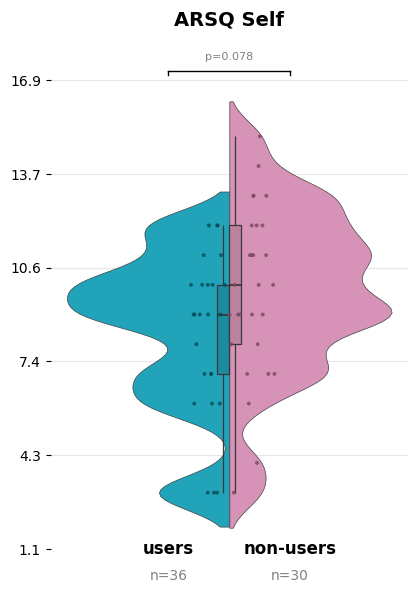 | 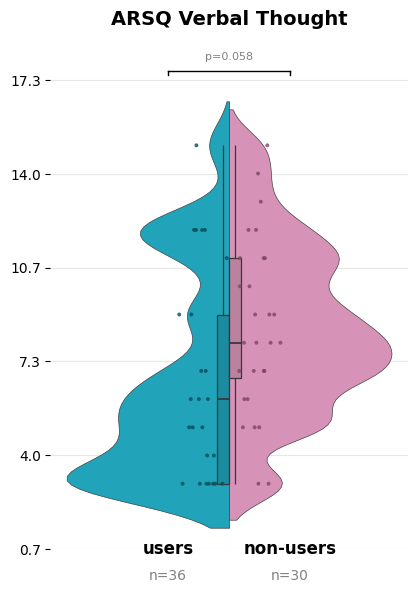 |
| --- | --- | --- | --- |
| 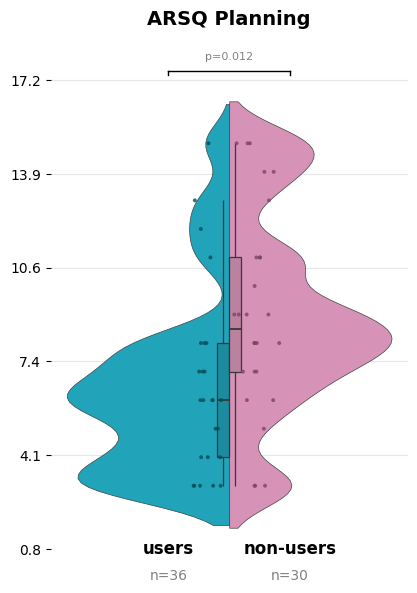 | 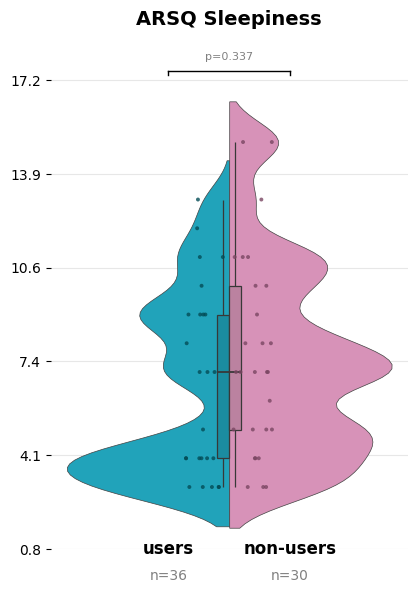 | 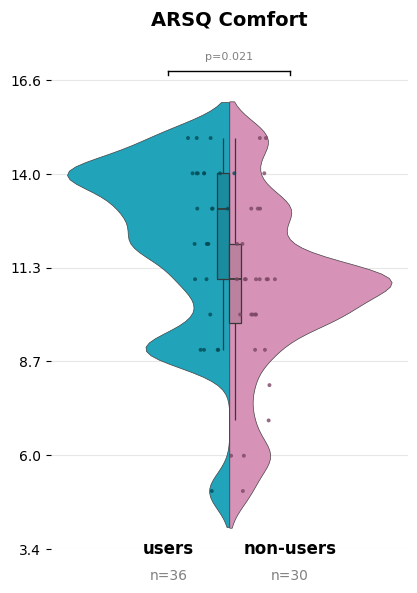 | 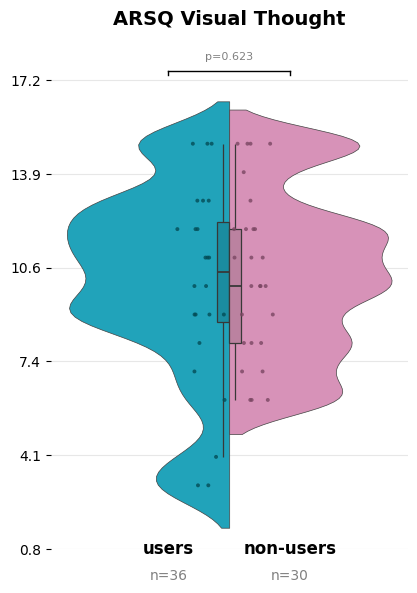 |
| 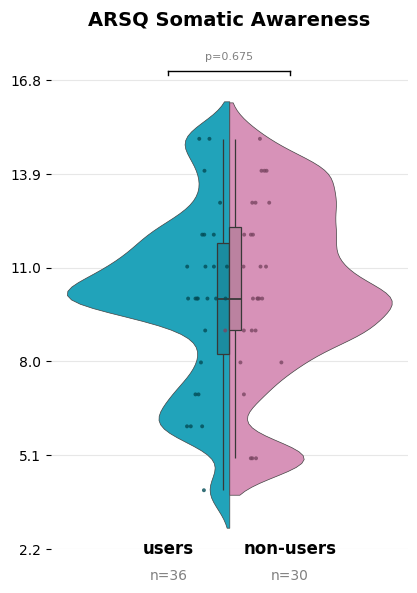 | 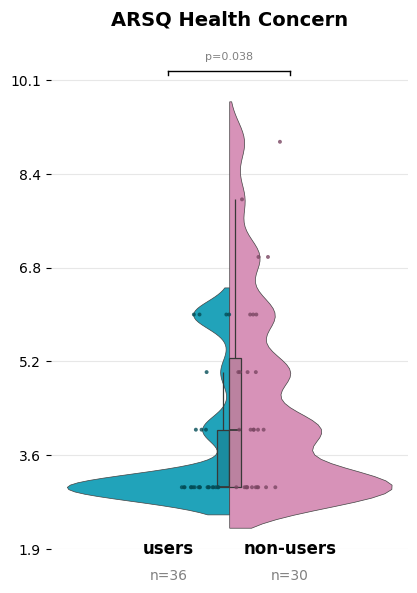 |  |  |

**Figure S8.** *Amsterdam Resting-State Questionnaire Subscale Distributions by Group (Dataset II).* Violin shapes represent probability density distributions; boxplots show median (center line), interquartile range (box), and whiskers (1.5× IQR). Individual data points shown as jittered dots (psychedelic users = blue/left; non-users = pink/right). Error bars represent range between first and third quartiles. ARSQ = Amsterdam Resting-State Questionnaire; *n* = sample size for each group.

### Supplementary Material 2

#### Preprocessing: oscillatory power and complexity

Custom MATLAB scripts based on EEGLAB functions (Delorme & Makeig, 2004) were used for automated preprocessing. Only electrodes common to both datasets (61 electrodes, 4 EOGs) were retained for each participant before preprocessing.

**Filtering and Resampling.** Signals were filtered with 1 Hz high-pass and 45 Hz low-pass FIR filters. Filter order was calculated using the formula *Pf* = 3.3 × *fs* / *Wt*, where *Pf* = filter order, *fs* = sampling frequency, and *Wt* = filter transition band width. Data were resampled to ensure consistency across datasets (Dataset I: 256 Hz; Dataset II: 250 Hz).

**Channel Rejection.** Bad channels were identified and removed using the NEAR plugin (criteria: flat signals >5 s; noisy channels >2 *SD* via periodogram analysis [1–45 Hz] 1-s windows with 66% overlap). An average of *M* = 2.29 (*SD* = 1.06) channels were removed per participant. Removed channels were reconstructed via spherical interpolation.

**Segmentation and Artifact Rejection.** Data were segmented into 4-s epochs. Epoch quality was assessed using three measures: (1) deviation of epoch's mean signal from channel-wise mean across epochs, (2) variance of signal within the epoch, and (3) maximum amplitude difference observed in epoch. Using the FASTER plugin for each measure, artifactual epochs were removed (thresholds: mean deviation, variance, max amplitude >2.5 SD. On average, *M* = 8.21 (*SD* = 2.31) epochs were removed per participant.

**Independent Component Analysis.** ICA with dimensionality reduction (36 principal components) identified artifact-related components via MARA (Winkler et al., 2011) resulting in *M* = 21.50 (*SD* = 4.60) components removed per participant.

**Quality Check and Referencing.** A final quality check removed channels with abnormally high power spectral density (>4 *SD* above mean; *M* = 0.82, *SD* = 4.76 channels removed) and interpolated them via spherical interpolation based on neighboring electrode activity. Signals were re-referenced to the average of both earlobe electrodes (A1, A2).

#### Preprocessing: effective connectivity

A more stringent (restrictive) preprocessing was employed, to minimize noise and ensure robustness of nDTF results. It should be noted that 54% of psychedelic neuroimaging studies fail contemporary standards for statistical correction and motion control (Linguiti et al., 2023).

**Dataset I.** Data underwent filtering within 2–47 Hz range utilizing windowed sinc linear phase FIR filters (high-pass order: 6758; low-pass order: 2552). Filter order was estimated using the formula *Pf* = 3.3 × *fs* / *Wt*, where *Pf* = filter order, *fs* = sampling frequency, and *Wt* = filter transition band width. Signals were then downsampled to 256 Hz, and virtual 4-s segmentation was employed for artifact rejection. Within the ASCT toolbox, bad channels were identified through the variance parameter using the IQR approach (threshold set to *Q*₁/*Q*₃ ± 7 IQR), calculated from EOG-corrected signals employing the recursive least square (RLS) method (Gomez-Herrero, 2007). Surviving original channels without EOG correction were re-referenced to the average of all EEG channels.

###### Dataset II. Signals underwent filtering within 2–47 Hz range (high-pass order: 3300; low-pass order: 1100) and were downsampled to 250 Hz. All other processing steps were identical to dataset I.

###### Both datasets. Trial-based artifact rejection was employed based on: variance (IQR threshold: 7), min/max difference (300 µV threshold), and muscle artifacts (elevated 35–47 Hz power). The remaining signals underwent decomposition using fastICA with dimensionality deflation for trial-based detection based on variance of IC time courses. ICA2 (24 iterations) with automated dual-stream classification employed two custom machine learning models (signal topographical patterns + spectral power; correlation between EOG channels and ICs). Final classification combined both models through weighted probability averaging to identify and remove ocular artifacts and extra-cerebral electrical interference. Decomposition allowed preservation of only brain-cortical components, facilitating more effective source-localization. Source reconstruction was conducted via minimum norm method (MNE; depth weighting *d* = 0.5), five-layer head model (SimBio toolbox; Vorwerk et al., 2018), SPM8 templates. ROI signals were reconstructed as sum of IC signals; scalar value represented by first principal component derived from three spatial vectors. MVAR model order was set to 7. Effective connectivity (nDTF) estimation used complete 4-s epochs. To limit the impact of spatial blurring of sources (leakage correction), symmetric multivariate orthogonalization was employed (ROInets toolbox; Colclough et al., 2015).

### Supplementary Material 3

#### ROI Specification

Default Mode Network (DMN) and Central Executive Network (CEN) regions were distinguished as following: (i) inferior parietal lobule areas—angular gyrus (ANG) assigned to DMN, supramarginal gyrus (SUP) assigned to CEN; (ii) inferior frontal gyrus (IFG) lateralization—left IFG assigned to DMN, right IFG assigned to CEN. Due to EEG spatial resolution limits, posterior cingulate cortex (PCC) and precuneus (PREC) were not individually specified and are combined as PREC/PCC.

###### Table S2. *MNI Coordinates for Regions of Interest in Default Mode, Central Executive, and Salience Networks.*

| **DMN** | | **CEN** | | **SN** | |
| --- | --- | --- | --- | --- | --- |
| **ROI** | **MNI**  (x y z) | **ROI** | **MNI**  (x y z) | **ROI** | **MNI**  (x y z) |
| L-IFG | -42 47 -7 | R-IFG | 48 47 -7 | L-dmACC | -6 6 38 |
| L-mPFC | -4 51 -7 | L-SFG | -15 20 62 | R-dmACC | 3 6 41 |
| R-mPFC | 5 53 -7 | R-SFG | 16 18 62 | L-INS | -37 17 -3 |
| L-aSTG | -59 -5 -6 | L-pSTG/TPJ | -66 -28 8 | R-INS | 37 18 -4 |
| R-aSTG | 59 0 -8 | R-pSTG/TPJ | 66 -30 11 |  |  |
| L-PREC/PCC | -9 -49 38 | L-ITG | -60 -56 -13 |  |  |
| R-PREC/PCC | 9 -49 38 | R-ITG | 63 -54 -13 |  |  |
| L-ANG | -51 -64 32 | L-SUP | -62 -35 36 |  |  |
| R-ANG | 54 -52 26 | R-SUP | 61 -40 41 |  |  |
| *Note.* MNI coordinates (x y z) represent the Montreal Neurological Institute’s stereotaxic space. L = left hemisphere; R = right hemisphere; ROI = Region of interest; DMN = Default Mode Network; CEN = Central Executive Network; SN = Salience Network; dmACC = dorsomedial anterior cingulate cortex; INS = insular cortex; IFG = inferior frontal gyrus; mPFC = medial prefrontal cortex; aSTG = anterior superior temporal gyrus; pSTG = posterior superior temporal gyrus; TPJ = temporoparietal junction; PREC = precuneus; PCC = posterior cingulate cortex; ANG = angular gyrus; SFG = superior frontal gyrus; ITG = inferior temporal gyrus; SUP = supramarginal gyrus. | | | | | |

### Supplementary Material 4

#### Additional results

| 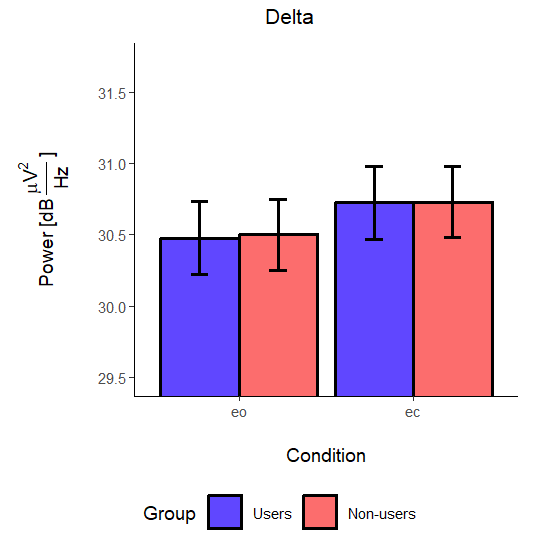 | 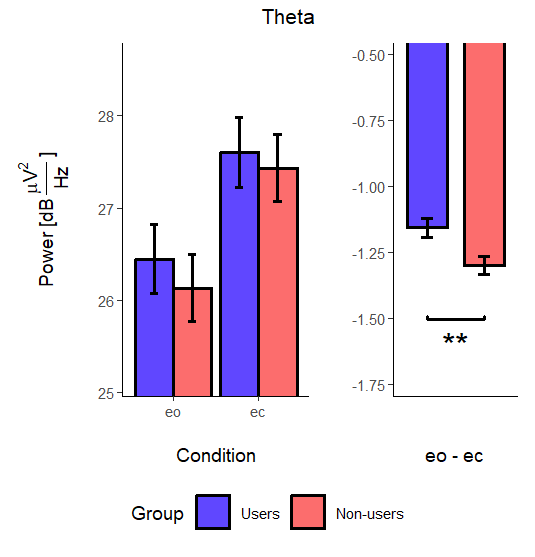 | 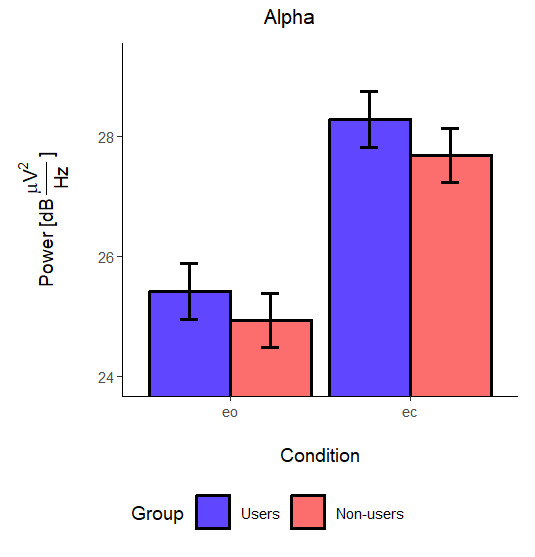 |
| --- | --- | --- |
| 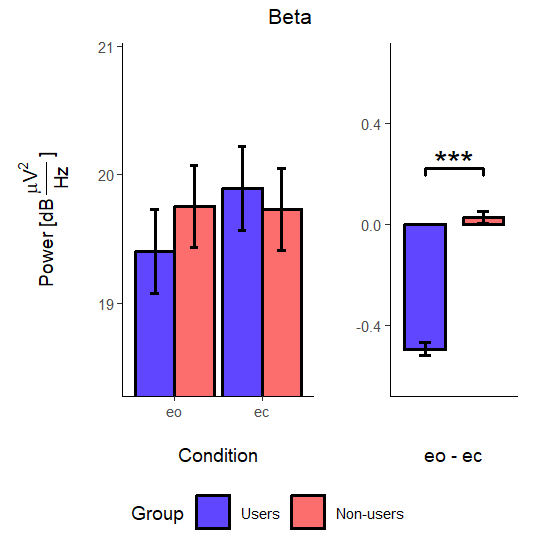 | 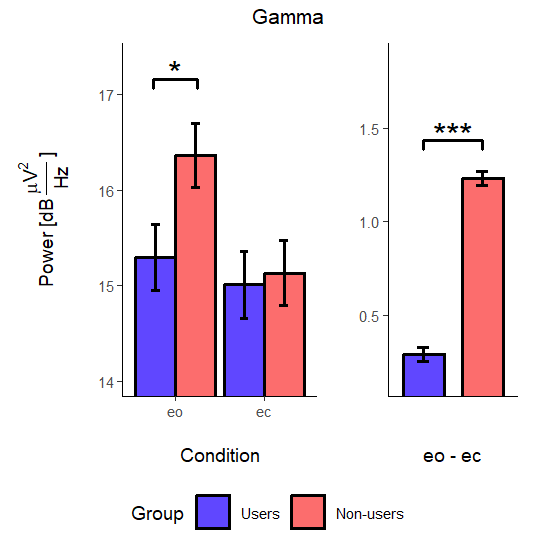 |  |

###### *Figure S9. Mean Power Spectral Density by Frequency Band, Condition, and Group. Error bars represent standard error.* Additional subplots for theta, beta, and gamma display between-group comparisons for the eyes-open minus eyes-closed contrast. *PSD =* power spectral density; dB µV²/Hz = decibels microvolts squared per hertz; eo = eyes-open condition; ec = eyes-closed condition. *p < .05. *p < .01. **p < .001.*.

**Table S3.** *Data Collection Site Effects in Oscillatory Power and Complexity.*

| **Measure** | **Direction** | **F(1, 110)** | ***p*** |
| --- | --- | --- | --- |
| **Oscillatory Power (PSD; dB μV²/Hz)** |  |  |  |
| delta | I > II | 13.37 | <.001*** |
| theta | n.s. | 0.46 | .499 |
| alpha | n.s. | 0.22 | .640 |
| beta | I > II | 7.74 | .006** |
| gamma | I > II | 30.69 | <.001*** |
| **Complexity (Lempel-Ziv)** | I > II | 4.95 | .028* |

*Note.* *F*-statistics from linear mixed-effects models. I = dataset I (Kraków); II = dataset II (Warszawa); PSD = power spectral density; dB µV²/Hz = decibels microvolts squared per hertz; n.s. — non-significant. *p* < .05. **p* < .01. ***p* < .001.

**Table S4.** *Regression Analysis Predicting Normalized U in Network-to-Network Directed Connections With Interaction Terms*

| **Parameters** | **Model 1** | **Model 2** | **Model 3** | **Model 4** |
| --- | --- | --- | --- | --- |
|  | *β* [95% CI] | *β* [95% CI] | *β* [95% CI] | *β* [95% CI] |
| **Covariates** |  |  |  |  |
| Intercept | — | — | −0.006 [−0.055, 0.042]  p = .808 | 0.012 [−0.037, 0.060]  p = .677 |
| Datasite (Warszawa) | — | — | — | −0.019 [−0.086, 0.049]  p = .635 |
| Condition (eyes-open) | — | — | −0.010 [−0.032, 0.011]  p = .388 | −0.009 [−0.031, 0.013]  p = .477 |
| **Frequency bands: main effects** (ref: delta) | |  |  |  |
| theta | — | — | 0.022 [−0.088, 0.132]  p = .732 | −0.000 [−0.046, 0.046]  p = .999 |
| alpha | — | — | 0.048 [−0.059, 0.156]  p = .408 | 0.036 [−0.042, 0.113]  p = .426 |
| beta | — | — | 0.030 [−0.065, 0.126]  p = .568 | 0.014 [−0.045, 0.074]  p = .682 |
| gamma | — | — | 0.015 [−0.071, 0.101]  p = .752 | 0.002 [−0.051, 0.055]  p = .951 |
| **Network Relations** (Model 1 only) | | |  |  |
| CEN→CEN | 0.012 [−0.035, 0.059]  p = .654 | — | — | — |
| CEN→DMN | 0.070* [0.003, 0.136]  p = .036 | — | — | — |
| CEN→SN | 0.078 [−0.018, 0.174]  p = .108 | — | — | — |
| DMN→CEN | −0.030 [−0.076, 0.016]  p = .230 | — | — | — |
| DMN→DMN | 0.026 [−0.009, 0.062]  p = .160 | — | — | — |
| DMN→SN | 0.032 [−0.037, 0.101]  p = .390 | — | — | — |
| SN→CEN | −0.023 [−0.103, 0.058]  p = .611 | — | — | — |
| SN→DMN | 0.016 [−0.047, 0.079]  p = .649 | — | — | — |
| SN→SN | 0.027 [−0.018, 0.071]  p = .263 | — | — | — |
| **Network × delta Interactions** (ref: CEN→CEN) | | |  |  |
| CEN→CEN | — | −0.011 [−0.062, 0.039]  p = .675 | ref | ref |
| CEN→DMN | — | 0.029 [−0.012, 0.070]  p = .189 | 0.040 [−0.003, 0.084]  p = .077 | 0.040 [0.003, 0.077]  p = .065 |
| CEN→SN | — | 0.057 [0.001, 0.112]  p = .053 | 0.068 [−0.016, 0.152]  p = .117 | 0.050 [−0.005, 0.105]  p = .125 |
| DMN→CEN | — | −0.053 [−0.128, 0.021]  p = .176 | −0.042 [−0.086, 0.002]  p = .070 | −0.037 [−0.077, 0.004]  p = .121 |
| DMN→DMN | — | 0.003 [−0.046, 0.052]  p = .905 | 0.015 [−0.016, 0.045]  p = .380 | 0.018 [−0.014, 0.050]  p = .330 |
| DMN→SN | — | 0.023 [−0.028, 0.073]  p = .412 | 0.034 [−0.034, 0.102]  p = .352 | 0.020 [−0.040, 0.080]  p = .558 |
| SN→CEN | — | −0.031 [−0.140, 0.078]  p = .601 | −0.020 [−0.091, 0.052]  p = .620 | −0.006 [−0.052, 0.039]  p = .817 |
| SN→DMN | — | −0.028 [−0.117, 0.061]  p = .566 | −0.016 [−0.070, 0.038]  p = .576 | −0.003 [−0.048, 0.042]  p = .912 |
| SN→SN | — | −0.005 [−0.076, 0.065]  p = .893 | 0.006 [−0.047, 0.059]  p = .828 | 0.005 [−0.049, 0.060]  p = .864 |
| **Network × theta Interactions** (ref: CEN→CEN) | | |  |  |
| CEN→CEN | — | 0.010 [−0.070, 0.091]  p = .822 | ref | ref |
| CEN→DMN | — | 0.066 [−0.026, 0.159]  p = .173 | 0.056* [0.014, 0.098]  p = .011 | 0.063* [0.019, 0.106]  p = .013 |
| CEN→SN | — | 0.083 [−0.034, 0.201]  p = .173 | 0.073* [0.004, 0.142]  p = .040 | 0.067 [0.003, 0.130]  p = .072 |
| DMN→CEN | — | −0.030 [−0.090, 0.030]  p = .362 | −0.040 [−0.093, 0.012]  p = .151 | −0.031 [−0.076, 0.014]  p = .234 |
| DMN→DMN | — | 0.014 [−0.044, 0.073]  p = .651 | 0.004 [−0.039, 0.047]  p = .876 | 0.013 [−0.029, 0.055]  p = .600 |
| DMN→SN | — | 0.036 [−0.055, 0.126]  p = .471 | 0.026 [−0.033, 0.084]  p = .418 | 0.035 [−0.034, 0.103]  p = .388 |
| SN→CEN | — | −0.016 [−0.087, 0.056]  p = .688 | −0.026 [−0.131, 0.079]  p = .653 | −0.004 [−0.055, 0.047]  p = .892 |
| SN→DMN | — | 0.021 [−0.041, 0.082]  p = .535 | 0.010 [−0.076, 0.097]  p = .827 | 0.027 [−0.026, 0.080]  p = .388 |
| SN→SN | — | 0.047 [−0.016, 0.110]  p = .168 | 0.037 [−0.040, 0.113]  p = .385 | 0.034 [−0.035, 0.102]  p = .405 |
| **Network × alpha Interactions** (ref: CEN→CEN) | | |  |  |
| CEN→CEN | — | 0.037 [−0.049, 0.124]  p = .435 | ref | ref |
| CEN→DMN | — | 0.105* [0.012, 0.199]  p = .030 | 0.068** [0.024, 0.112]  p = .003 | 0.068** [0.026, 0.110]  p = .006 |
| CEN→SN | — | 0.093 [−0.011, 0.197]  p = .079 | 0.056 [−0.015, 0.127]  p = .136 | 0.040 [−0.019, 0.100]  p = .240 |
| DMN→CEN | — | 0.016 [−0.060, 0.093]  p = .699 | −0.021 [−0.071, 0.030]  p = .454 | −0.015 [−0.056, 0.026]  p = .541 |
| DMN→DMN | — | 0.067 [−0.004, 0.138]  p = .076 | 0.030 [−0.024, 0.084]  p = .308 | 0.036 [−0.015, 0.087]  p = .232 |
| DMN→SN | — | 0.061 [−0.018, 0.140]  p = .147 | 0.024 [−0.043, 0.091]  p = .508 | 0.027 [−0.044, 0.098]  p = .526 |
| SN→CEN | — | −0.007 [−0.109, 0.095]  p = .903 | −0.044 [−0.155, 0.067]  p = .470 | −0.021 [−0.075, 0.034]  p = .524 |
| SN→DMN | — | 0.051 [−0.039, 0.141]  p = .295 | 0.014 [−0.086, 0.114]  p = .801 | 0.037 [−0.028, 0.102]  p = .323 |
| SN→SN | — | 0.026 [−0.046, 0.099]  p = .509 | −0.011 [−0.102, 0.081]  p = .836 | −0.002 [−0.071, 0.067]  p = .966 |
| **Network × beta Interactions** (ref: CEN→CEN) | | |  |  |
| CEN→CEN | — | 0.019 [−0.049, 0.086]  p = .614 | ref | ref |
| CEN→DMN | — | 0.088* [0.004, 0.173]  p = .041 | 0.070** [0.021, 0.118]  p = .005 | 0.075** [0.031, 0.118]  p = .003 |
| CEN→SN | — | 0.097 [−0.016, 0.210]  p = .092 | 0.078 [−0.008, 0.164]  p = .081 | 0.066 [−0.005, 0.137]  p = .111 |
| DMN→CEN | — | −0.017 [−0.077, 0.043]  p = .593 | −0.036 [−0.090, 0.018]  p = .216 | −0.027 [−0.076, 0.021]  p = .335 |
| DMN→DMN | — | 0.032 [−0.018, 0.083]  p = .239 | 0.013 [−0.032, 0.058]  p = .589 | 0.029 [−0.015, 0.073]  p = .259 |
| DMN→SN | — | 0.027 [−0.053, 0.108]  p = .537 | 0.009 [−0.059, 0.076]  p = .813 | 0.014 [−0.061, 0.089]  p = .756 |
| SN→CEN | — | −0.004 [−0.093, 0.086]  p = .941 | −0.022 [−0.134, 0.089]  p = .720 | −0.004 [−0.061, 0.052]  p = .893 |
| SN→DMN | — | 0.038 [−0.032, 0.108]  p = .320 | 0.019 [−0.072, 0.110]  p = .696 | 0.034 [−0.026, 0.094]  p = .330 |
| SN→SN | — | 0.043 [−0.026, 0.111]  p = .253 | 0.024 [−0.062, 0.110]  p = .616 | 0.012 [−0.069, 0.093]  p = .807 |
| **Network × gamma Interactions** (ref: CEN→CEN) | | |  |  |
| CEN→CEN | — | 0.004 [−0.051, 0.059]  p = .900 | ref | ref |
| CEN→DMN | — | 0.059 [−0.021, 0.138]  p = .161 | 0.055* [0.008, 0.102]  p = .025 | 0.054* [0.013, 0.096]  p = .024 |
| CEN→SN | — | 0.059 [−0.065, 0.183]  p = .379 | 0.055 [−0.038, 0.148]  p = .269 | 0.035 [−0.030, 0.101]  p = .358 |
| DMN→CEN | — | −0.067* [−0.121, −0.014]  p = .020 | −0.071* [−0.126, −0.016]  p = .012 | −0.062* [−0.109, −0.016]  p = .021 |
| DMN→DMN | — | 0.016 [−0.026, 0.057]  p = .489 | 0.012 [−0.023, 0.046]  p = .536 | 0.024 [−0.012, 0.059]  p = .246 |
| DMN→SN | — | 0.012 [−0.083, 0.108]  p = .820 | 0.009 [−0.064, 0.081]  p = .829 | 0.006 [−0.067, 0.080]  p = .888 |
| SN→CEN | — | −0.056 [−0.135, 0.023]  p = .188 | −0.060 [−0.162, 0.042]  p = .280 | −0.041 [−0.099, 0.017]  p = .222 |
| SN→DMN | — | −0.002 [−0.064, 0.060]  p = .958 | −0.006 [−0.087, 0.075]  p = .894 | 0.009 [−0.053, 0.071]  p = .806 |
| SN→SN | — | 0.024 [−0.035, 0.082]  p = .456 | 0.020 [−0.037, 0.076]  p = .518 | 0.017 [−0.041, 0.074]  p = .614 |
| **Model statistics** | | |  |  |
| ***R²*** | .115 | .162 | .164 | .057 |
| ***F*** | 74.78 | 20.03 | 19.90 | 12.05 |
| ***p*** | .132 | .195 | .197 | .305 |

*Note.* Model 1: contrast ~ network_relation. Model 2: contrast ~ network_relation × bands. Model 3: contrast ~ city + eyes + network_relation × bands (merged datasets I+II). Model 4: contrast ~ city + eyes + network_relation × bands (separate datasets). Higher *β* indicates higher connectivity in the psychedelic users group. All *p*-values, standard errors, and confidence intervals are permutation-based (10,000 permutations). CI = confidence intervals; CEN = Central Executive Network; DMN = Default Mode Network; SN = Salience Network; ref = reference category. **p* < .05. ***p* < .01.

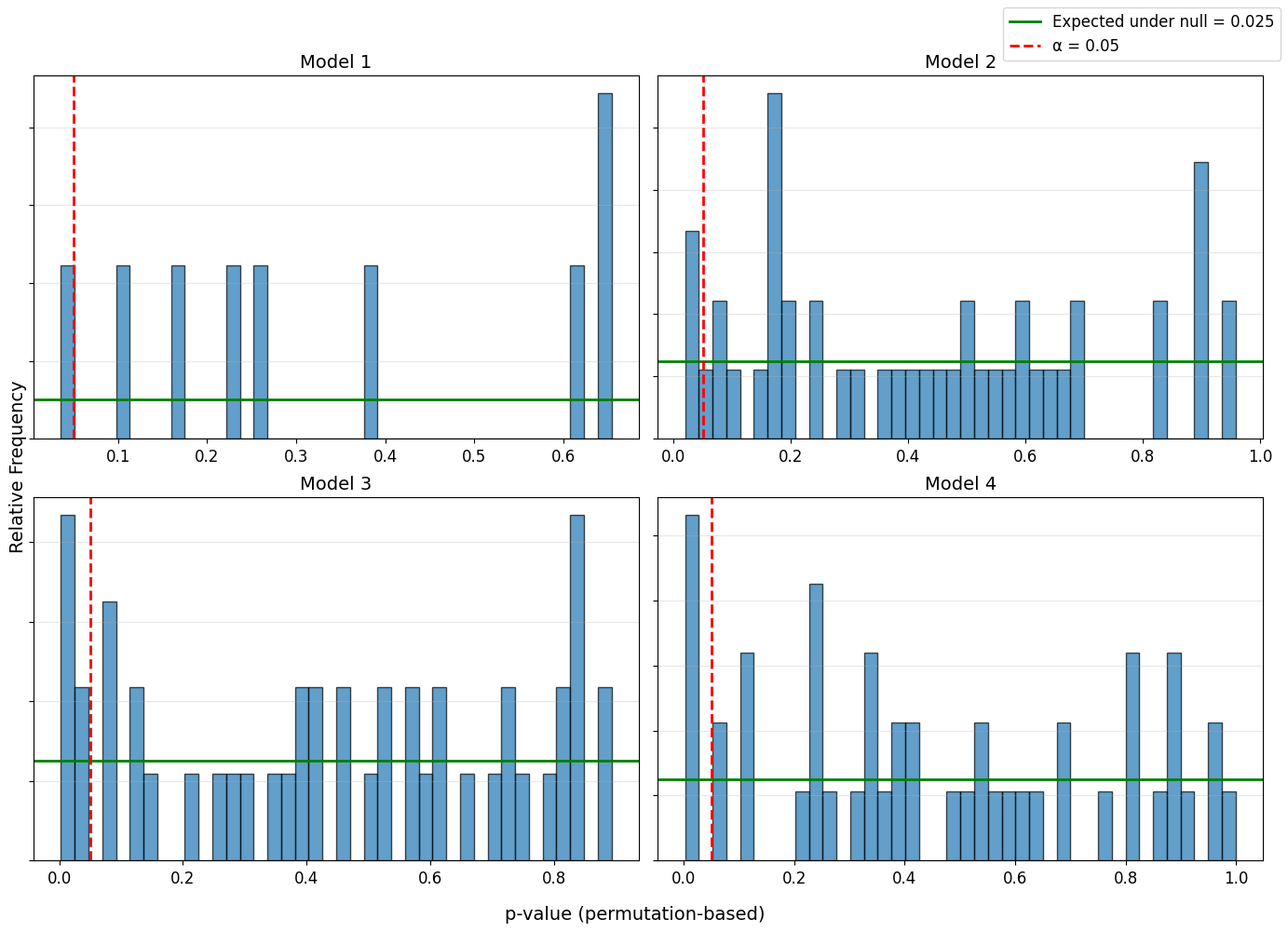

**Figure S10.** *Distribution of Permutation-Based p-Values Across All Four Regression Models.* Histograms display frequency distributions of permutation-based *p*-values for all regression coefficients in each model. Red dashed vertical line indicates α = .025; green horizontal line indicates expected frequency under null hypothesis (α = 0.025). Models 1–4 correspond to analyses in Table S4.

**END OF SUPPLEMENTARY MATERIALS**
